# Time-varying interactions change coexistence mechanisms

**DOI:** 10.1101/2025.11.28.690940

**Authors:** Violeta Calleja-Solanas, Ignasi Bartomeus, Oscar Godoy

## Abstract

Species interactions are fundamental to maintaining diversity. However, it remains unclear how environmental variability modifies the structure, sign, and strength of species interactions that ultimately affect coexistence mechanisms. Here, we combined structural stability theory with time-varying population models to study the temporal variation of niche and fitness differences over nine years of annual plants in Mediterranean grasslands. Rainfall variation modulated species-specific responses in performance, and changed the magnitude of self-limitation and the variability in the strength of inter-specific interactions for both competition and facilitation. These changes implied substantial variation in yearly niche and fitness differences, which ultimately impacted community stability. Additional simulations show that this full biological reorganization due to rainfall variation can maintain diversity by promoting temporal shifts in winners and losers compared to a single climatic year. Our results highlight the need to incorporate time-varying interactions to understand species coexistence under contrasting environmental conditions.

## 1 Introduction

A century of research on biotic interactions has shown that they are not fixed species attributes. Their sign and strength can shift substantially with changes in climate, community composition, and local conditions (Ushio et al., 2018; García-Callejas et al., 2021; Buche et al., 2025), leading to major alterations in the overall structure of species interactions (CaraDonna, Petry, et al., 2017; Poisot et al., 2015). Although this context dependency is widely recognized (Chamberlain et al., 2014; Pearce-Higgins et al., 2015; Liu and Gaines, 2022; Catford et al., 2022), it has rarely been incorporated into population models that describe the temporal dynamics of ecological communities under natural conditions (Godoy, Soler-Toscano, et al., 2024; Nguyen et al., 2025). As a result, we still have a limited understanding of how temporal variation in the network of biotic interactions relates to changes in the stability of ecological communities and the underlying mechanisms that promote species coexistence.

To address this limitation in a context of environmental variation, it is essential to recognize that species persist in an ever-changing world (Godoy, Soler-Toscano, et al., 2024; Chesson, 2017; Vicente-Serrano et al., 2025), yet such variability is often simplified. Logistical constraints are a major reason. Manipulative experiments measuring changes in species interactions, defined as the average percapita effect of one species on another, are extremely labor-intensive when conducted across more than a few climatic scenarios (Van Dyke et al., 2022; Wainwright, HilleRisLambers, et al., 2019; Matías et al., 2018; Muehleisen et al., 2025). Even when changes in community composition are predicted, they are typically examined at the species-pair level (Ploughe et al., 2019). Such a level of species richness focuses on direct interactions while overlooking the indirect effects that interaction chains exert on maintaining diverse communities (Levine, Bascompte, et al., 2017). Most studies also concentrate on a single taxon or guild, resulting in a limited understanding of how temporally dynamic responses compare across groups with distinct evolutionary histories, such as plants and insects. Additional challenges arise from theoretical simplifications. For example, climate-driven predictions of community composition generally emphasize competitive interactions (Hallett et al., 2019; Van Dyke et al., 2022), neglecting the widespread role of facilitation (CaraDonna, Burkle, et al., 2021; Buche et al., 2025; Losapio et al., 2021) and the fact that shifts from competition to facilitation frequently occur under increasing climatic stress (Bertness and R. Callaway, 1994; Maestre et al., 2009).

A framework suited to explore the temporal variation in coexistence mechanisms for multispecies communities that include both facilitation and competition is the theory of structural stability. This theory, combined with population models, presents a toolbox to identify how species interactions define the range of variation in species’ performance compatible with the feasibility of an ecological community (i.e., all species persist in the long-term with positive abundances) (Saavedra et al., 2017; Medeiros et al., 2021; Allen-Perkins et al., 2023; Song et al., 2020). Analogous to other theories, structural stability defines two coexistence mechanisms. Structural niche differences (Saavedra et al., 2017) represent the extent of opportunities for species to coexist (Godoy, Bartomeus, et al., 2018; Allen-Perkins et al., 2023), and are assessed by computing the size and the shape of the so-called feasibility domain. This domain is determined by the interactions and contains the range of potential intrinsic growth rates for which all species have a feasible equilibrium. Conversely, structural fitness differences define asymmetries in species’ performance (Saavedra et al., 2017). Structural stability predicts that species can coexist when the species’ realized growth rates remain within this domain. Conversely, if the species’ realized growth rates are outside, some species will decrease their population in the short-term, and they will eventually be excluded (Medeiros et al., 2021; Allen-Perkins et al., 2023). Although structural stability was not originally explored to include time-varying model parameters (Rohr et al., 2014; Saavedra et al., 2017), there is no theoretical limitation to include them (Godoy, Soler-Toscano, et al., 2024). Our main theoretical expectation is that including environmental variation will change species’ intrinsic growth rates and their interactions, which in turn will temporally affect structural stability through changes in coexistence mechanisms. In particular, we predict that less structurally stable communities occur in years when environmental variation reduces niche differences and increases fitness differences. Ultimately, we hypothesize that, if sustained, the loss of stability will lead to the local extinction of specific species, thereby impoverishing local communities.

Here, we illustrate how temporal variation in environmental conditions drives changes in coexistence mechanisms. In our case study, rainfall drives these changes by affecting species’ intrinsic growth rates and their interactions within the community. We focus on rainfall because prior experimental work has shown that it shapes diversity and community composition through its effects on species’ vital rates (Dore, 2005; Pearce-Higgins et al., 2015; Knapp et al., 2002; Thibault and Brown, 2008; Sandel et al., 2010) and on species interactions (Matías et al., 2018; Wainwright, HilleRisLambers, et al., 2019; Van Dyke et al., 2022; Muehleisen et al., 2025). To do so, we draw on a highly replicated multispecies ecological community followed under natural conditions. We recorded changes in species abundances for 7 annual plants in grasslands over 9 years and across 324 plots. The system’s main environmental variable, rainfall *θ*_*t*_, ranged from years with typical rainfall to multiyear droughts, with anomalies of roughly 300 mm below the 85-year average rainfall ⟨*θ*⟩. We parameterized a Lotka–Volterra-like model in which intrinsic growth rates, 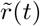, and interaction coefficients, *Ã*(*t*), vary through time in response to rainfall, Fig. 1a–c; Eq. (2), while also incorporating temporal variation in spatial stochasticity in 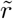. We fit a time-varying model using Bayesian inference, allowing us to explicitly quantify uncertainty by estimating posterior distributions for each parameter. This also means that interactions quantify the effect of the abundance of species *j* on the per capita growth of species *i* (Abrams, 1987; Novak et al., 2016), and then go beyond just encoding co-occurrence (Yin and Rudolf, 2024). Although this approach requires rich multi-annual data to estimate interactions with confidence, thereby restricting the number of species to consider, it has the important advantage that we do not impose the sign of species parameters. We can explore whether including time-varying species parameters and stochasticity improves the description of abundance evolution. Then, examining patterns of both positive and negative interactions sheds light on the ongoing debate over whether environmental variation alters species interactions in predictable ways (Maestre et al., 2009) or whether changes are largely context-dependent (Catford et al., 2022; Chamberlain et al., 2014), with randomness emerging from demographic stochasticity, spatial heterogeneity, unmeasured environmental factors, or simply sheer luck (Shoemaker et al., 2020).

**Figure 1:**
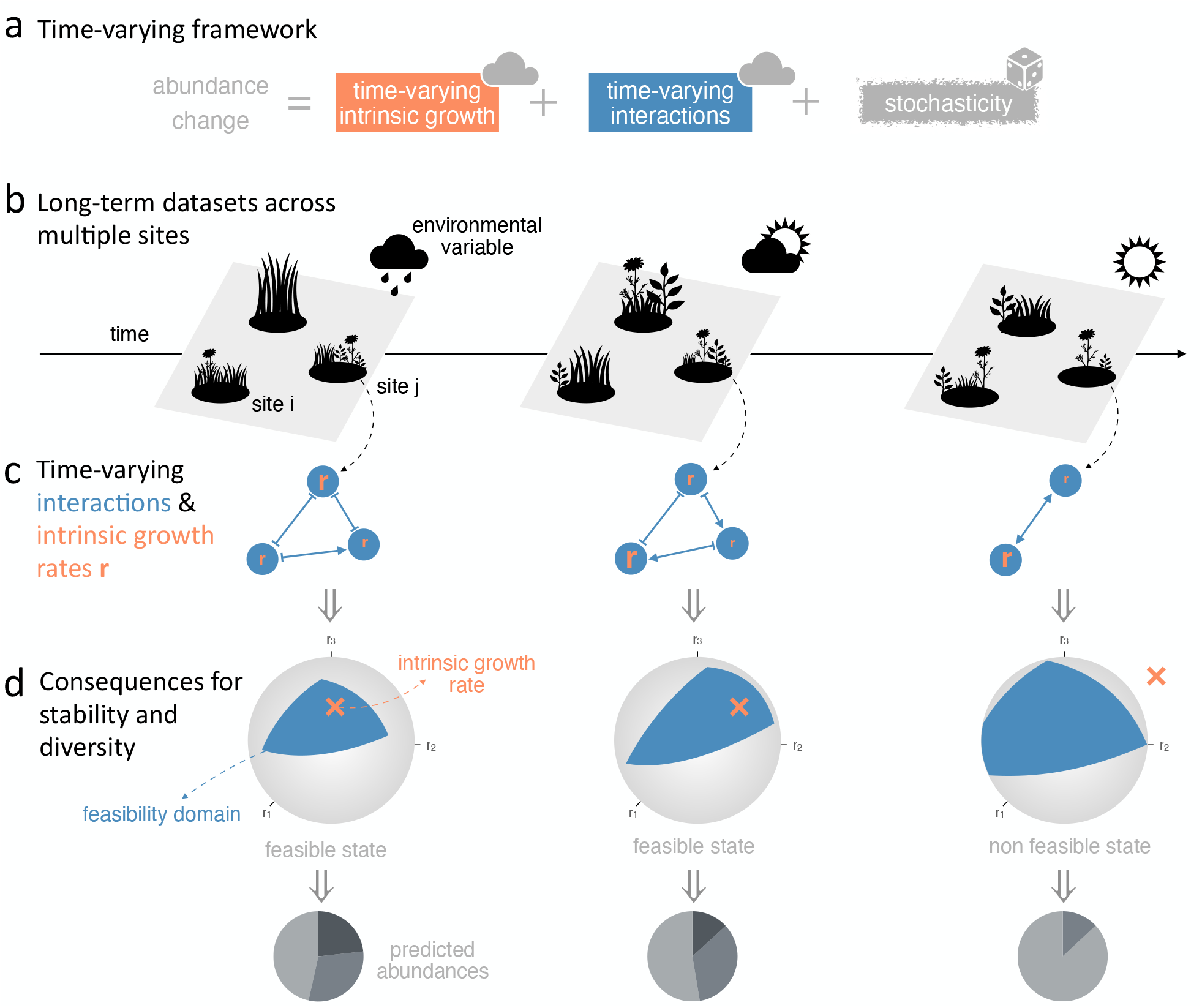
Illustration of the theoretical framework in a toy community of 3 species. (a) Estimation of interactions and intrinsic growth rates that vary through time according to changes in environmental factors. For each dataset, (b) thanks to well-resolved empirical observations from multiple sites constrained to rainfall, we can disentangle the time-varying and stochastic contributions of species’ parameters to community dynamics, (c) obtaining time-varying species interactions *Ã* (represented as blue networks) and intrinsic growth rates 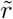 for each site. (d) In turn, these changes in species parameters modify the emerging properties of ecological communities, such as their stability (represented as the position of 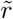, orange cross, within the feasibility domain, blue shape) and diversity.

## 2 Material and Methods

### 2.1 Empirical data

Time series containing data on species populations are from a field study replicated across multiple areas. It is located at Caracoles Ranch, an annual grassland of 2680 ha at Doñana National Park, southwest Spain (García-Callejas et al., 2021). The data gathered during *>* 360 hours of empirical observations associated with this study contains 9 years (2015 - 2023) of field surveys of local neighborhoods and species abundances following a spatially explicit design of 9 sites (8.5 m *×* 8.5 m along a 1 km *×* 200 m area) dominated by annual plant species with no perennial species present, Fig. S1. In addition, each site was divided into 36 subplots of 1 m *×* 1 m, with 0.5 m-wide aisles, for a total of 324 subplots and 20.412 data points. Soil conditions at these sites vary due to a salinity gradient and nutrient content. For our analysis, we considered the *n* = 7 plant species present every year. Therefore, there is sufficient statistical power to incorporate the impact of time-varying environmental factors.

Thanks to this unique dataset, we obtained empirical estimates of intrinsic growth rates 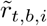 that are variable in time *t* (across 9 years) plus a stochastic spatial effect across the different sampling sites *b*, and interactions *Ã*_*t,ij*_variable in time. The species belong to disparate taxonomic families and have different functional profiles (Table 1), but are primarily subjected to the same environmental driver, that is, annual rainfall. Moreover, they have recently experienced drought periods (Fig. S2), making them suitable for studying the consequences of droughts on community structure. The anomalies to study the effect of rainfall on stability and community composition are calculated as the difference from the long-term 85-year average (520 mm).

### 2.2 Time-varying framework

Our theoretical approach is based on the Ricker model, a discrete version of the generalized Lotka-Volterra equations (Levine and HilleRisLambers, 2009). In its classic, deterministic form, changes in the abundance of species *i* at time 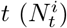 are described by:

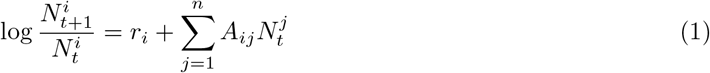

where *r*_*i*_ is the intrinsic growth rate of species *i* and *A* is the interaction matrix of the *n* species. This baseline model does not depend on any temporal variable. The Ricker model is commonly applied to plant communities (Mayfield and Stouffer, 2017) because it can naturally accommodate both positive and negative interactions with varying strength, something that other standard population models such as the Beverton-Holt cannot do.

To explicitly link community dynamics to environmental fluctuations, we introduced time-varying species responses in intrinsic growth rates 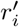 and interactions *B*_*ij*_ via effects of environmental conditions at each time *θ*_*t*_. In our system, the primary environmental driver is rainfall *θ*_*t*_. Furthermore, to account for local contingencies across our different sampling sites *b*, we incorporated hierarchical, site-level stochastic effects on species performances, *u*_*i,b*_. Our proposed time-varying framework reads:

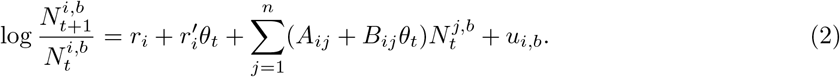

The time-varying terms of Eq. (2) involve a deterministic term on the growth rates 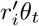 and interactions *B*_*ij*_*θ*_*t*_. By fitting this full model via Bayesian inference, we obtained joint posterior distributions for all baseline parameters and their environmental modifiers. From these posteriors, we calculated the distributions of the realized interaction matrices (*Ã*_*t,ij*_) and realized intrinsic growth rates 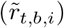 for any given rainfall condition and site:

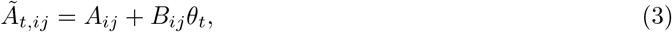

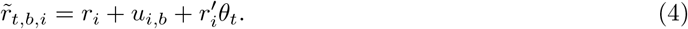

Hence, species interactions change annually in response to an environmental variable, and species exhibit different performance depending on that environmental driver and the site *b* where they are sampled.

To study the long-term behavior of the system, in which the average per capita growth rates are all zero, we time-averaged the multispecies Ricker model. Assuming the populations persist over a sufficiently long time, the abundance change (left-hand side of Eq. (2)) tends to zero after some transient:

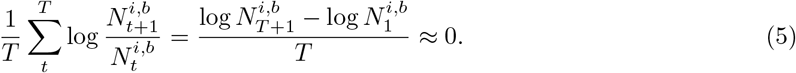

Applying the time-averaging operator 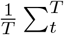, and denoting the time average of a variable *x*_*t*_ as 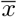, we can map into the classic linear form 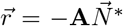, by defining the effective time-averaged interaction matrix elements 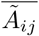, Eq. (6), and the effective time-averaged intrinsic growth rates, Eq. (7):

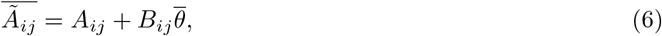

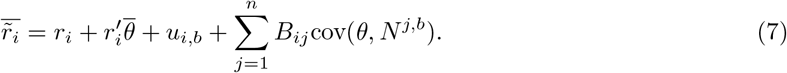

The covariance term appears only in Eq. (7). It accounts for the effect of averaging a non-linear function over the temporal sequences of environmental conditions *{θ*_1_, *θ*_2_, …, *θ*_*t*_, …, *θ*_*T*_ *}* and population sizes *N*. As a consequence, two different sequences of environmental conditions with the same mean *θ*, such as a constant environment (*θ*_1_ = *θ*_2_ = *…* = *θ*_*T*_, ∀*t*) and an oscillating one (*{θ*_1_, *θ*_2_, *θ*_1_, *θ*_2_, … *}*), have the same 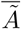, but differ in their covariance and then in their 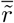 (Supplementary Information, SI 1).

### 2.3 Bayesian Modeling Approach

Using Bayesian inference, we parameterized the community model with our multi-year empirical dataset. This approach allowed us to estimate the sign and strength of interactions, as well as intrinsic growth rates, as posterior probability distributions rather than fixed-point estimates (SI 2). Nevertheless, to test the robustness of our results to the choice of inference method, we also estimated the parameters using the R ‘nlme’ package version 3.1.164 for generalized linear mixed-effects models (SI 3), obtaining parameter values close to the means of the posterior distributions (Figs. S11– 16).

To evaluate the predictive performance of the time-varying framework, Eq.(2), we fit models of increasing complexity (SI 2.1) sharing the same priors, and evaluated their expected log predictive density using leave-one-out cross-validation, via the R packages ‘brms’ version 2.22.0 and ‘loo’ version 2.8.0.

### 2.4 Structural stability analysis

To connect community stability with rainfall-driven changes in species’ intrinsic growth rates 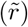 and in their interaction structure (*Ã*), we applied structural stability theory. It pinpoints the set of species’ performances for which communities are feasible. While environmental conditions are traditionally assumed to modify only species’ performance (Rohr et al., 2014), we extended this approach here by (1) incorporating environmentally driven variation also in interactions and (2) making that dependence explicit (Eqs. 3 and 4). In doing so, structural stability links the range of environmental conditions compatible with the persistence of multiple interacting species (i.e., the coexistence opportunities) to the structure of species interactions (Saavedra et al., 2017). The larger these coexistence opportunities, measured as the size of the feasibility domain Ω(*Ã*), the larger the range of species’ intrinsic growth rates the community tolerates without losing species, Fig. S3. Because our inferred interactions encompass both positive and negative signs, and intrinsic growth rates can also be negative, we generalized the standard calculation of Ω(*Ã*). We normalized the solid angle that determines the feasibility domain, i.e., the (hyper)area of the convex hull defined by the column vectors of *Ã*, by the area of the whole unit (hyper)sphere, rather than by the area defined by all the vectors with only positive elements. In this way, we could accommodate our situation, in which interactions are also positive and intrinsic growth rates can be negative (Fig. 1d). A graphical representation of these concepts is in Fig. S3.

In a time-varying framework, both the shape and size of the feasibility domain may vary since they are dictated by *Ã* (Allen-Perkins et al., 2023; Grilli et al., 2017), which may reorganize with rainfall. The position of 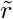 may also change over time, as shown in Eq. (4). Because we estimated these parameters using Bayesian inference, we fully propagated this parameter uncertainty by calculating all structural stability metrics across the posterior draws. For the time-averaged approach, the effective time-averaged interaction matrix 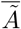 only depends on the mean environmental conditions 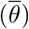, so two different environmental trajectories will have the same feasibility domain (size and shape) if they have a common mean environment 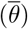. On the contrary, time-averaged intrinsic growths 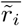 not only depend on 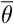 but also on a fluctuation-dependent term, the covariance between rainfall and the time series of species abundances. This covariance encodes the differences that arise from two different environmental trajectories. Then, time-varying environmental conditions do not change coexistence opportunities, but can alter the intrinsic growth rate vector across the feasibility domain via the covariance term.

The structural approach maps conceptually onto coexistence mechanisms of the Modern Coexistence Theory (Godoy, Bartomeus, et al., 2018). The structural analog of niche differences is precisely the size of the feasibility domain Ω(*Ã*) since it quantifies coexistence opportunities (Saavedra et al., 2017). Similarly, the structural analog of fitness differences is the angular deviation (in radians) between a realized vector of intrinsic growth rates and the incenter of the feasibility domain, which maximizes the possibility of a feasible solution (Allen-Perkins et al., 2023). To identify which species are more vulnerable to extinction under greater fitness differences, we computed the slope between rainfall anomalies and each species’ distance to its exclusion boundary. (Medeiros et al., 2021; Allen-Perkins et al., 2023; Lepori et al., 2024). The feasibility domain is the set of directions of 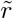 for which all species attain positive equilibrium abundances; it is determined entirely by the interactions, being the cone spanned by the columns of *Ã*, and its borders are the directions at which one species’ abundance reaches zero (Fig. S3). Its size Ω(*Ã*) is therefore the structural analog of niche differences (Saavedra et al., 2017), because it measures how much variation in performance the community tolerates before losing a species. For each rainfall condition and draw, the system may lie in a different position within a feasibility domain. The position of 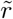 is the structural analog of fitness differences: the incenter is the direction equidistant from all borders, so the angular deviation between 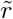 and the incenter quantifies the asymmetry in species’ performances (Allen-Perkins et al., 2023). Coexistence then requires this angular deviation to remain within the extent of the domain, the structural counterpart of requiring fitness differences to be smaller than niche differences. We accordingly quantify structural stability as the distance between 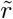 and the nearest border of the feasibility domain (Medeiros et al., 2021; Allen-Perkins et al., 2023; Lepori et al., 2024). This measure increases with the size of the domain and decreases with the angular deviation of 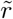, so a community can lose stability either because its interactions shrink the domain or because asymmetries in performance push 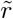 towards a border. We adopted a sign convention in the distance to the borders: it is negative when 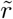 is outside the feasibility domain. These generalized analyses were implemented by contributing to the R package anisoFun v2.0 https://github.com/RadicalCommEcol/anisoFun.

### 2.5 Network measures of temporal variability

We computed several properties of the interaction network’s structure and tracked how they varied with extreme environmental conditions. In particular, we tested the extent to which interactions were conserved over time, measuring the temporality of the networks between two times *m* and *l* (Bauzá Mingueza et al., 2023): 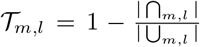, where | ⋂_*m,l*_| is the size of the intersection (the common interactions of the two networks), whereas | ⋃_*m,l*_ | is the size of the union (the interactions that appear at least at one of the times). Consequently, temporality scales from 0 (identical network topology) to 1 (complete turnover of interactions). This metric follows the same foundational concepts as beta diversity; while beta diversity describes the turnover of multiple species in space, *T*_*m,l*_ describes the turnover of interactions over time.

To calculate temporality, we binarized the realized interaction matrices within each posterior draw, so that parameter uncertainty propagates into the credible interval of the metric rather than into the definition of the network. For each draw and year *t*, we defined an interaction as present in a given year if its strength exceeded half the standard deviation of the off-diagonal elements of that same matrix |*Ã*_*t,ij*_| ≥ *σ*_*t*_*/k*, with *k* = 2. Using this criterion, we calculated the temporality of the unsigned networks. We also computed signed temporality by analyzing the turnover of positive and negative unweighted interactions separately.

To characterize the overall quantitative structure, we measured the diagonal dominance across the posterior draws. For each draw, we calculated the percentage of species satisfying the condition: 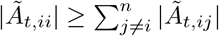, and tracked how it varied with environmental conditions. When the absolute diagonal element *Ã*_*t,ii*_ is larger than the sum of the absolute off-diagonal elements *Ã*_*t,ij*_ for all species, the interaction matrix is diagonally dominant, meaning that species limit themselves more than they are limited by other species. While classical studies often focus on global differences between *competitive* interactions to infer niche differentiation (resulting in greater values)(Chesson, 2000), examining positive and negative interactions allows us to discover more opportunities for species to coexist (Godoy, Calleja-Solanas, et al., 2026). Theory predicts that diagonally dominant matrices stabilize population dynamics and maintain diversity (Saavedra et al., 2017), a coexistence mechanism empirically supported across diverse ecological communities (Adler et al., 2018; Daniel et al., 2024; Calleja-Solanas et al., 2025).

### 2.6 Simulations

We numerically explored how different environmental conditions affect the coexistence of the communities by accounting for different rainfall scenarios with the same mean 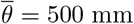: constant rainfall similar to the 85-year average, *θ*_cte_ = 500 mm, decreasing rainfall *θ*_↓_ from 600 to 400 mm, increasing rainfall *θ*_↑_ from 400 to 600 mm, a triangular wave *θ*_∧_ alternating between 600 and 400 mm, and a sinusoidal sequence *θ*_∼_ centered at 500 mm with 100 mm of amplitude and a 5-year period. To run these simulations, we drew 9000 parameter values from our posterior distributions and parameterized the multispecies Ricker equations for each draw until they reached (quasi)stationarity. We then recorded which parameter combinations led to all species persisting. We performed these simulations to obtain a computational approximation of the size of the feasibility domain. Then, we calculated the time-averaged 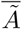 and 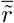 (neglecting the transient), Eqs. (6) and (7), to compare their feasibility predictions with the simulations. Since we have inferred the parameters of the time-varying model, Eq. (2), we can also simulate how biodiversity unfolds in the short term, during our 9 year sampling period, under hypothetical rainfall regimes, by producing abundance time series for a given rainfall sequence *θ* = *{θ*_1_, …, *θ*_*T*_ *}*, such as: constant rainfall equal to the 85-year average ⟨*θ*⟩ = 520 mm, oscillations between the minimum, min(*θ*) = 225 mm, and maximum, max(*θ*) = 625.5 mm, recorded rainfalls in our study period, increasing rainfall from ⟨*θ*⟩ to max(*θ*) + *σ*(*θ*) ≃ 700 mm, and decreasing rainfall and drought from ⟨*θ*⟩ to min(*θ*) − *σ*(*θ*) ≃ 100mm. To fully propagate the parameter uncertainty, we generated an ensemble of simulations for each scenario by sampling from the joint posterior distributions. The simulations were run for the same duration as our empirical dataset (9 timesteps). To track biodiversity changes, we calculated the Shannon index at each timestep *t* and site *b* across the simulation ensembles: 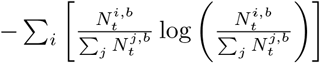 Ultimately, these simulations have three possible outcomes: (i) the presence of a persistent group of species (yielding a positive Shannon diversity), (ii) the complete species extinction of the community, or (iii) the dominance by a single species (in the last two cases, Shannon diversity is zero).

## 3 Results

The time-varying model best predicted observed plant abundance dynamics (Fig. S4.). The explanatory power was satisfactory according to the leave-one-out validation, and the time-varying model outperformed simpler models, such as those considering only time-varying intrinsic growth rates or no time-variability, based on the standard error of component-wise differences (Tables 2 and 3). With the predictive power of the time-varying framework well established, we examined the posterior distributions of interactions and intrinsic growth rates. We found that their central tendencies and credible intervals shift in both strength and sign across the observed temporal gradient of rainfall variation (SI 4).

### 3.1 Rainfall variability produces contrasting effects on the temporal networks’ structure

To describe the overall structure of the variation in interactions due to rainfall changes, we computed the temporality *T*_*t,t*+1_, a Jaccard-inspired metric that measures how strongly interaction networks differ from one year to another. Larger rainfall changes were positively associated with larger differences between interaction networks (Fig. 2a), and temporality also exhibited comparable trends when calculated using only facilitative or only competitive interactions (Fig. S5). Results were robust to the value of the scaling constant *k* (Fig. S6).

**Figure 2:**
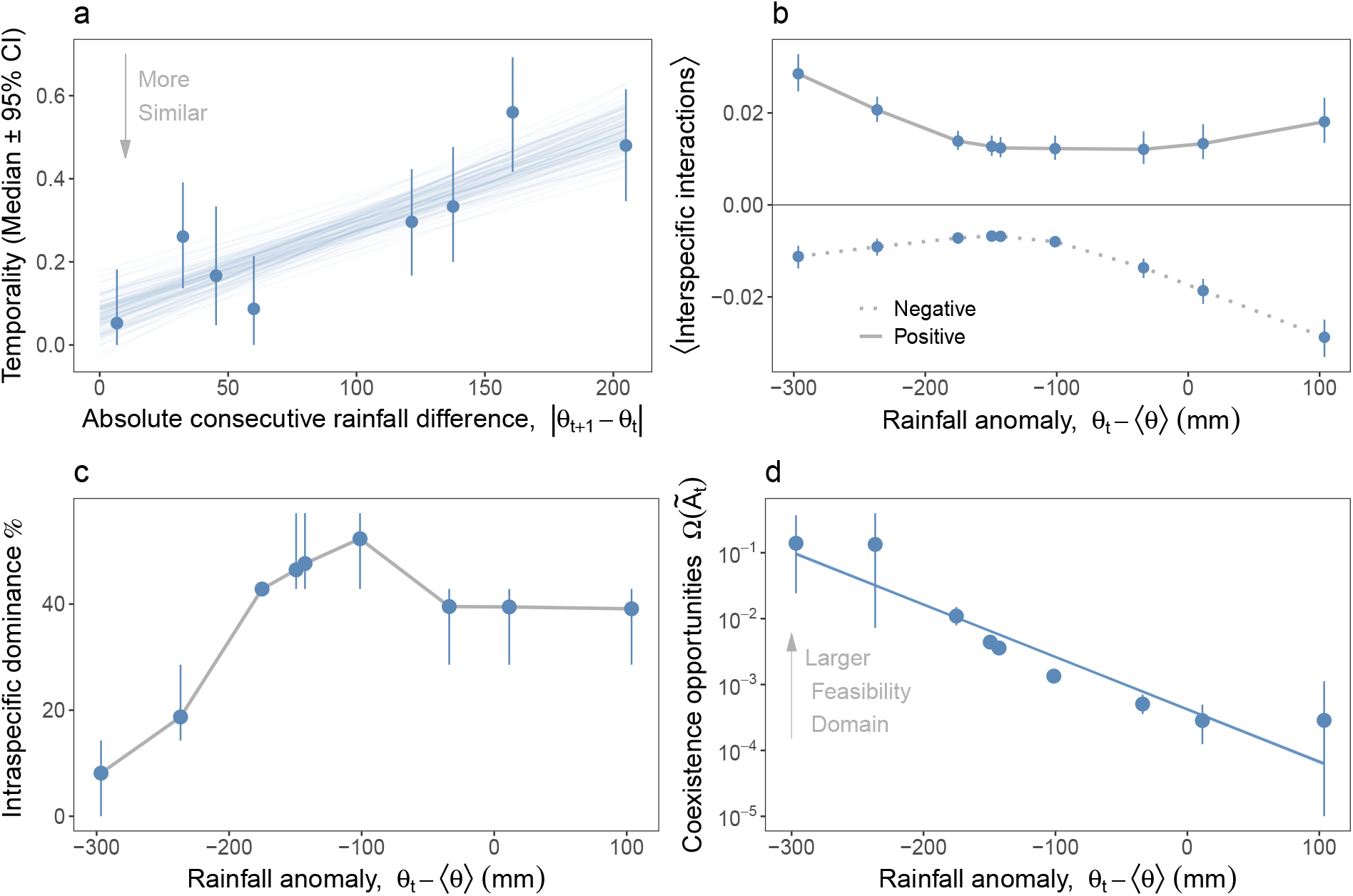
Structural changes in interaction networks over time. **a** Temporality, the degree of turnover of two consecutive interaction matrices. Blue solid lines are least-squares fits obtained separately within each of 100 randomly chosen posterior draws, so that their spread reflects posterior uncertainty in the interactions. **b** Mean interspecific positive (solid) and negative (dotted) interactions over species versus rainfall anomalies. **c** Changes in the percentage of species whose self-regulation dominates their interspecific interactions with rainfall anomalies. Smaller values indicate less self-regulation. Gray lines for visualization guidance. **d** Coexistence opportunities (size of the feasibility domain for each realized interaction matrix Ω(*Ã*_*t*_)) with respect to rainfall anomalies. For each rainfall anomaly, properties of the interaction network are calculated with 1000 posterior draws. Solid blue lines are Bayesian generalized linear model fits for 100 draws. Note that gray lines in **b** and **c** are for visualization purposes. SI 4 contains the posterior distributions and temporal evolution of the values of the time-varying networks.

Crucially, the predominant sign of these networks shifted with rainfall anomalies, defined as deviations from the long-term seasonal average. Facilitation was more common in drier years, while competition dominated in wetter years (Fig. 2b). Near the long-term mean ⟨*θ*⟩, posterior distributions clustered around zero, indicating predominantly weak interactions. However, under the driest and wettest anomalies, credible intervals widened and shifted away from zero; that is, interactions became stronger. This switch, in which harsh environments induce a transition from competitive to facilitative interactions, is highly consistent with the stress gradient hypothesis.

To evaluate the balance between self-limitation and interspecific competition, we quantified annual diagonal dominance of the drawn networks. The diagonal dominance declined with increasing drought (Fig. 2b), indicating a weakening of self-limitation relative to interspecific interactions. As self-limitation becomes relatively weaker with increasing rainfall anomalies in drier years, we would expect a reduction in coexistence opportunities, as measured by the size of the feasibility domain. Yet, we found the opposite. The drier the year, the larger the opportunities to coexist (estimated by the size of the feasibility domain Ω(*Ã*_*t*_), Fig. 2c). In the driest year (−297mm relative to the average), coexistence opportunities increased.

### 3.2 Rainfall variability shapes species performance, determining winners and losers

Interannual changes in rainfall affected the time-varying intrinsic growth rate 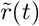 in predictable ways. Intrinsic growth rates generally increased with positive rainfall anomalies (wetter years) (Fig. 3a). These differences in how rainfall drives species’ intrinsic growth rates promoted fitness differences (the distance between the intrinsic growth rates of all species and the vector for which extinctions were less probable, the incenter of the feasibility domain): drier years increased fitness differences for the plant community (Fig. 3b).

**Figure 3:**
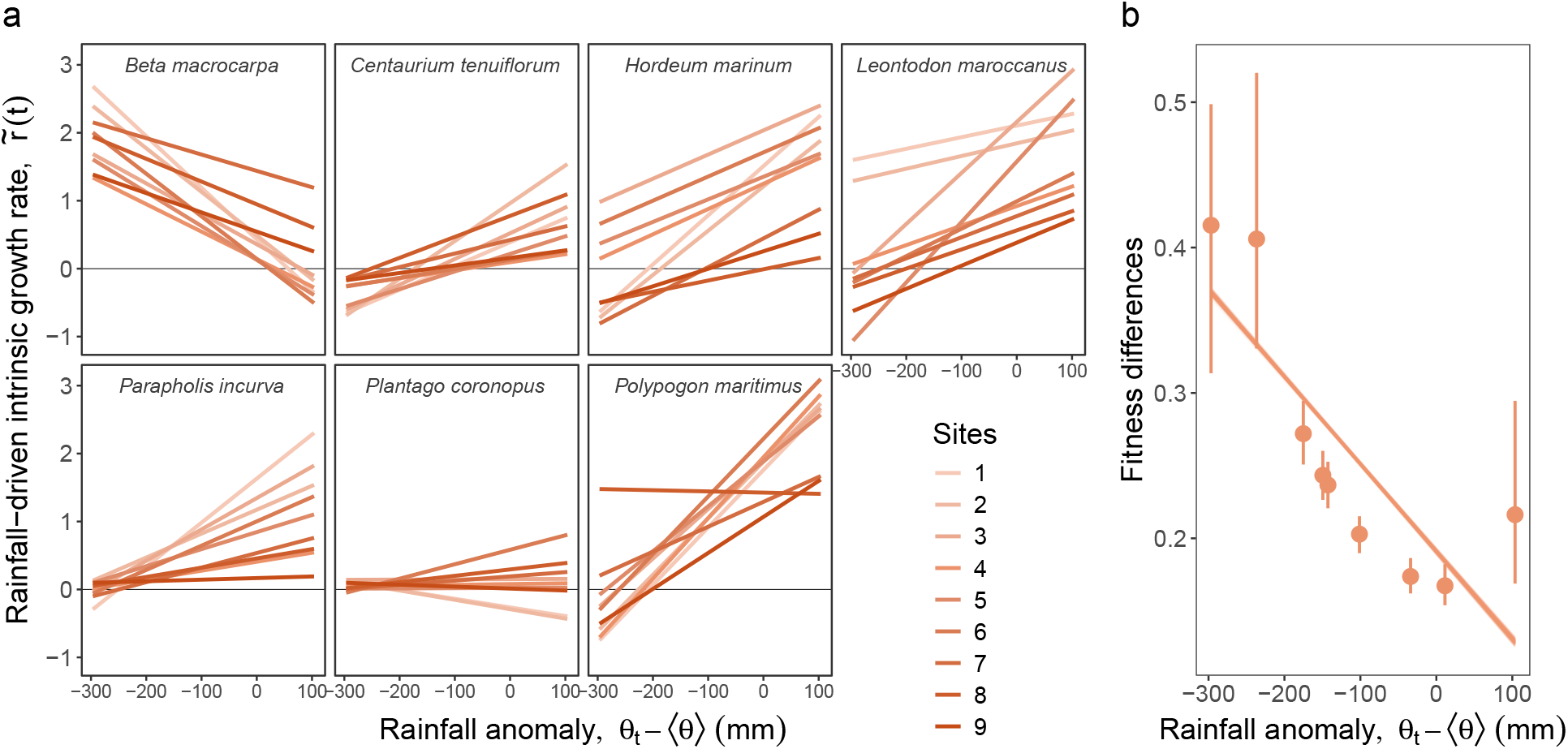
Species’ performance explained by deterministic effects of rainfall and spatial context dependency. **a** Realized intrinsic growth rates for all sampled sites. A relationship parallel to the x-axis indicates no sensitivity to rainfall anomalies, and a greater slope variation for a species means more sensitivity to the site’s conditions. Only the posterior distribution mean is plotted here. **b** Fitness differences associated with rainfall anomalies, measured as the distance between 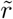 and the incenter. Each point is the average across sites of 1000 posteriors draws, and solid orange lines are Bayesian generalized linear model fits for 100 draws. SI 4 contains the temporal evolution of the values of the time-varying 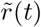.

### 3.3 Assessing structural stability under time-varying interactions and intrinsic growth rates

Building on our earlier results, we examined how structural stability emerges from the interplay between time-varying coexistence opportunities (Fig. 2c) and differences in species’ intrinsic growth rates (Fig. 3b). That is, the balance between niche and fitness differences. We found that rainfall anomalies can strengthen or weaken structural stability depending on the direction of the anomaly, Fig. 4a, and species. Furthermore, some sites even show negative values in the extreme anomalies, indicating likely extinctions in the long-term. For most species, the distance to their exclusion border decreased as rainfall increased (negative slopes in Fig. 4b), so these species sat farther from extinction in drier years. *Parapholis incurva* is a counterexample, for which both intrinsic growth rate and distance to the border increase with rainfall. The wide error bars in Fig. 4a show that, for extreme anomalies, some sites maintained positive structural stability, helping to preserve diversity, whereas others are near the stability boundary, risking species loss.

**Figure 4:**
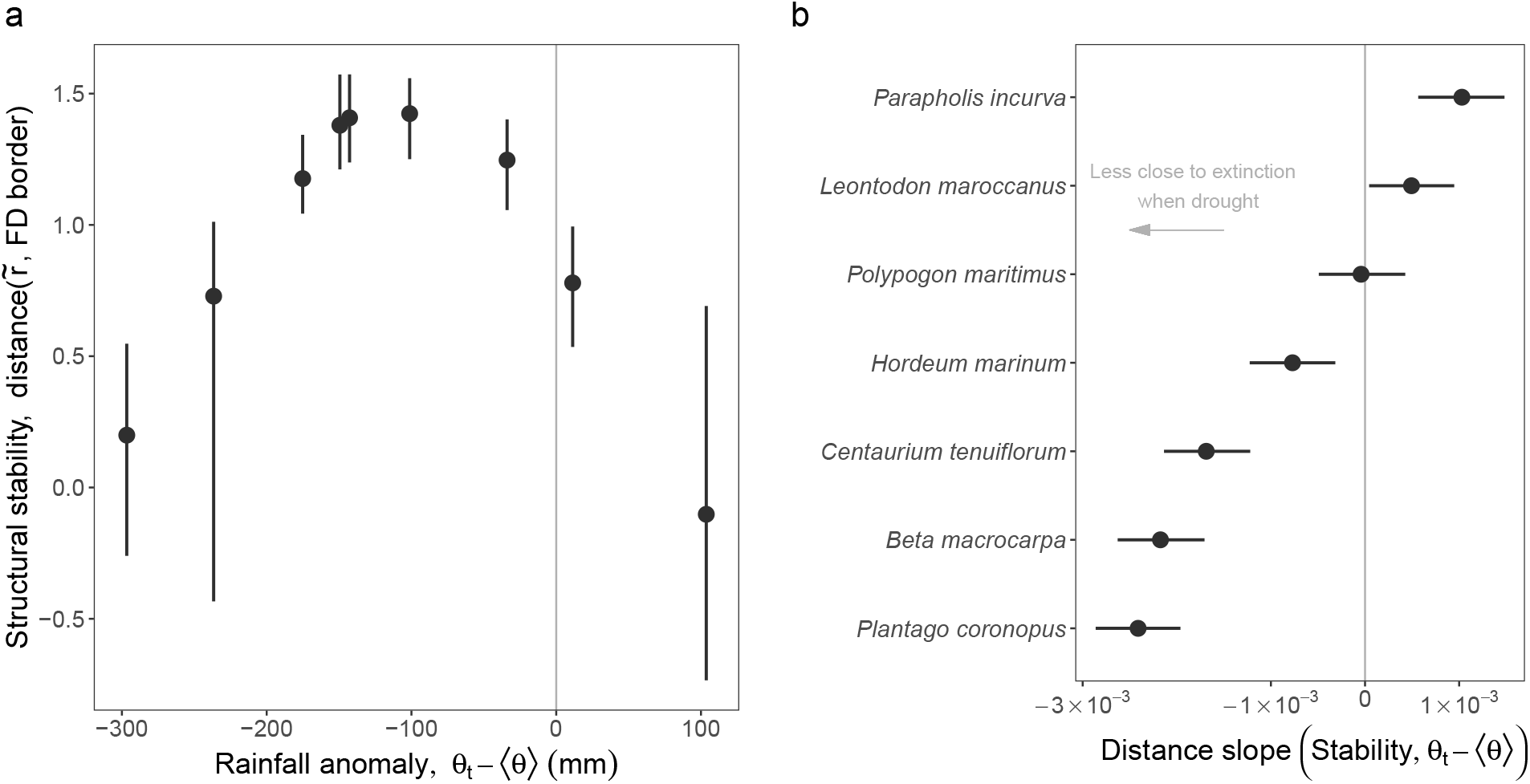
Consequences on stability of temporal changes in coexistence mechanisms. **a** Structural stability as the average over the whole community and sites of the species distance between intrinsic growth rates and the nearest border of the feasibility domain (FD) where a species goes extinct. **b** Slopes of the distance between intrinsic growth rates and the nearest border of the feasibility domain (FD) for each species as a function of rainfall anomalies. The trends for each species are in Fig. S7. In our sign convention, positive distances denote that a species is more prone to persist in wetter years, and negative slopes indicate greater species persistence in drier years. Error bars represent the standard deviations across all sites for 1000 draws each.

### 3.4 Predictions of biodiversity loss and the amplification of spatial context dependence

An aspect worth highlighting of our framework is that it also predicts how biodiversity will change under different rainfall regimes. We started by exploring the 9-year short term but with constant rainfall, equal to the historical average ⟨*θ*⟩. Although real environmental conditions fluctuate continuously, this baseline provides a useful reference for comparing other rainfall regimes. We then explored three additional regimes: a gradual decrease in rainfall, a gradual increase, and an oscillation between minimum and maximum values. For each regime, we simulated population dynamics using the parameterized model for the corresponding time-varying rainfall levels and calculated the Shannon diversity index annually (Fig. 5a).

**Figure 5:**
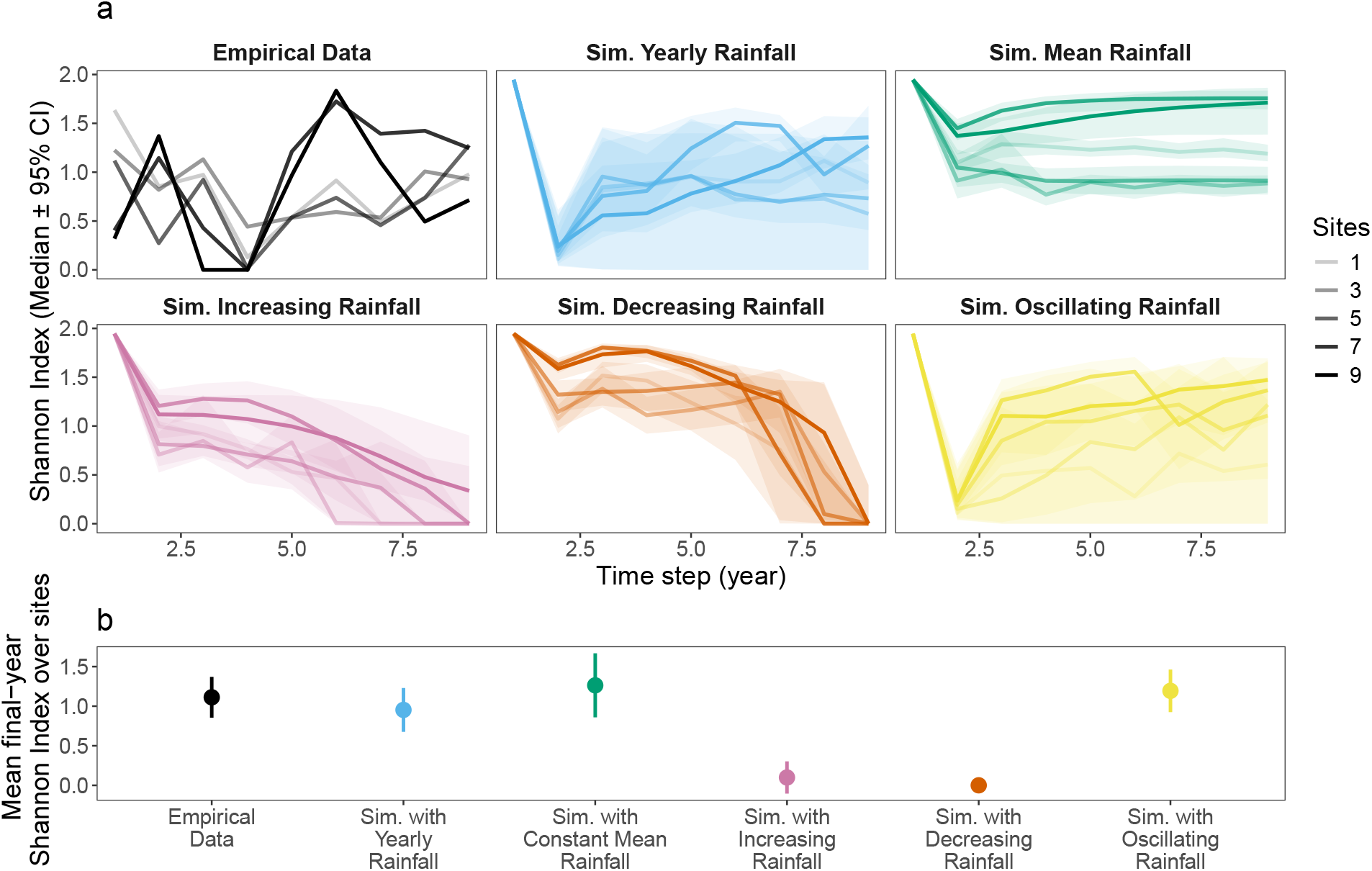
Consequences on diversity of rainfall patterns. **a** Shannon diversity index for different simulated rainfall regimes. Besides the index for the empirical abundances (black), the lines show 1000 simulations from our estimated model for different scenarios: rainfall varies according to the recorded values, rainfall does not vary and is equal to the average, increases over time, decreases, and oscillates between the maximum and minimum values over 9 years. **b** Shannon indices for the final state of the simulations. Since our community size is *n* = 7, the maximum value of the index is 2.

For most sites in the study systems, we found that the constant, increasing, and oscillating rainfall regimes have a positive Shannon diversity at the end of the simulations (Fig. 5b), consistent with the empirical dataset. Note that the documented rainfall does not change gradually but instead shows peaks and troughs—despite the recent drought trend (Fig. 2), a pattern resembling the oscillating regime. This oscillating scenario also produces the highest Shannon diversity in the plant system. In contrast, the gradual drought regime generally causes a collapse of Shannon diversity to zero, leaving at most a single species in the community.

Simulations accounting for long-term dynamics with a *θ*_∧_ scenario (triangular fluctuations in rainfall) obtained the most feasible communities at the end of our simulations (1762*/*9000 ≈ 20%) than any other scenario, followed by constant rainfall *θ*_cte_ (1534 ≈ 17%) and sinusoid fluctuations in rainfall *θ*_∼_ (1471 ≈ 16%). The decreasing and increasing rainfall scenarios had only 881 (∼ 1%) and 234 (∼ 0.3%) feasible communities, respectively. This means that the oscillating scenarios have among the largest numerically calculated feasibility domains. Interestingly, from these feasible communities, the time-averaged intrinsic growth rate overestimates and underestimates the feasibility of some scenarios depending on the sites (Fig. S8).

## 4 Discussion

Our results provide rigorous estimations, combining theory, population models, and empirical data, of how ecological communities respond to temporal changes in environmental conditions (rainfall variation in our study). The theoretical framework applied here also shows how these responses translate to coexistence mechanisms and the emerging structural stability. Taken together, the results tell a single story in three steps. First, environmental conditions reorganize the interaction network, with drier years weakening self-limitation and shifting the balance of interspecific interactions towards facilitation. Such facilitation enlarges the feasibility domain and thus increases coexistence opportunities. Second, environmental conditions also reorganize species’ performances asymmetrically because species differ in how strongly they benefit from water availability. Drier years generate larger fitness differences, with site-level stochasticity amplifying or dampening these differences across the landscape. Since these two coexistence mechanisms respond to drought in opposite directions, their combination results in the highest values of structural stability occurring at precipitation levels close to the long-term average. Finally, and consistent with this second finding, our simulations show that the community can sustain the highest diversity when rainfall remains close to its long-term average. Our system, however, is far from static. Rainfall has strongly fluctuated over the nine years of study, and our simulation shows that such fluctuations are the alternative scenario in which the observed diversity of the system can also be maintained. Simulating other scenarios, such as directional trends or multi-year drought, result in species extinction and drive a collapse of the system. We explain each of these three main results in greater detail below.

We find that the decrease in self-limitation coupled with the increase in facilitation explained the larger opportunities for coexistence assessed in drier years. The increase in facilitation and its connections with niche differences agree in general terms with the stress gradient hypothesis (Maestre et al., 2009; CaraDonna, Petry, et al., 2017), and the role of facilitation in enlarging species’ niche (Bruno et al., 2003). Moreover, recent theoretical advances indicate that, in the case of a reduction of self-limitation, facilitation between species can alternatively promote coexistence opportunities by preventing the extinction of poor-performing species that persist thanks to the presence of others (Godoy, Calleja-Solanas, et al., 2026; Rohr et al., 2014). Therefore, our results suggest that, in years of low rainfall, our case study could potentially withstand a broader range of intrinsic growth rates thanks to the prevalence of positive interspecific interactions (Gross et al., 2024). However, it is essential to compare the magnitude of coexistence opportunities (i.e., niche differences) with the differences in species’ intrinsic growth rates (i.e., fitness differences) under each rainfall scenario for this statement to hold.

The temporal variation in intrinsic growth rates was asymmetric, consistent with previous studies showing that species do not benefit equally from environmental conditions, leading to winners and losers under changing environmental regimes (Van Dyke et al., 2022; Wainwright, HilleRisLambers, et al., 2019; Thibault and Brown, 2008; Knapp et al., 2002; Matías et al., 2018). This contrast reflects fundamental ecological differences. Plants rely differently on water availability for germination, growth, and reproduction (Wainwright, Staples, et al., 2018; Knapp et al., 2002). These heterogeneous performances collectively result in plants performing better and more similarly in wetter years. A direct prediction from this finding is that the annual plant community is less prone to species loss over wet years. The underlying mechanism explaining this decrease in fitness differences is that the more similar the performance across species, the greater the likelihood of coexistence because the community is more centrally located within the feasibility domain, and farther from its exclusion borders (Saavedra et al., 2017; Allen-Perkins et al., 2023; Nguyen et al., 2025). Moreover, intrinsic growth rates do not depend solely on rainfall. Site-level stochasticity (spatial heterogeneity) modulates the realized intrinsic growth rates 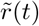, either amplifying or dampening rainfall effects. Consequently, some sites buffered poor years while others intensified negative impacts. Future work should test whether certain taxa are inherently more sensitive to stochasticity in their responses to rainfall variation. However, whether the reduction in fitness differences we observe in wetter years translates into greater levels of coexistence depends on how it combines with the opposite trend that rainfall has on promoting coexistence opportunities.

Contrary to our initial expectations, the two mechanisms promoting structural stability respond to drought in opposite directions. Although coexistence opportunities increased under lower rainfall, asymmetries in intrinsic growth rates 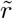 also became more variable, producing a nonlinear effect on structural stability at the community level. Although species showed clear preferences for dry or wet conditions in the absence of interactions and in their distances to the exclusion border, the overall structural stability is highest under conditions close to the long-term rainfall average values and not at the rainfall extremes. A paradigmatic example is *Centaurium teniflorum*. Despite obtaining higher intrinsic growth rates in wetter years, it slightly approximates the extinction border in wet years. This discrepancy between direct and realized responses indicates that interaction networks strongly reshape species’ climatic niches. While the role of biotic interactions in modifying climatic niches has been discussed at biogeographic scales (Wiens, 2011; Stephan et al., 2021; Galiana et al., 2023), it has rarely been evaluated locally. Our results show that the temporal reorganization of interaction networks and performances prevents species from capitalizing on extreme rainfall conditions. Instead, most species persist best under rainfall near the long-term mean.

From a methodological perspective, while standard structural stability analysis evaluates the feasibility of a fixed-point equilibrium (*N*_*t*+1_ = *N*_*t*_), in natural systems, such fixed points are rarely, if ever, attained. Instead, by calculating the feasibility domain Ω(*Ã*_*t*_) for each environmental condition, we are quantifying yearly, the instantaneous attractors of the community. The theoretical equilibrium defined by the realized parameters of a given year dictates the immediate directional pull on population trajectories during that time step. If a species’ realized intrinsic growth rate 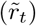 falls outside Ω(*Ã*_*t*_), the model exerts an instantaneous pressure to extinction for that species (Allen-Perkins et al., 2023). Structural stability is then a proxy for stability, indicating where the community would go if not altered by changes in environmental conditions. In fact, even though some rainfall conditions set low coexistence opportunities, the fact that the parameters are changing may prevent that fixed point with low coexistence opportunities, or even unfeasible, from being materialized. In the study of these non-equilibrium dynamics, the long-term coexistence depends not only on the magnitude of species’ environmental responses, but also on the specific temporal sequence of those changes. Because our mechanistic framework explicitly models both, it is suited to test how different temporal regimes unfold over time.

We tested the effect of temporal variation on coexistence by simulating the community under contrasting rainfall sequences, tracking how the community’s position relative to the feasibility domain shifted from year to year. Coexistence, indicated by a final positive Shannon diversity index, emerged under three scenarios, each driven by a distinct stabilizing mechanism. First, simulations replicating the observed rainfall sequence successfully maintained community diversity. Because these species naturally coexist in our empirical dataset, this outcome provides support for our Bayesian estimates of species’ intrinsic growth rates and their interactions. Second, coexistence was also maintained under the constant, average rainfall scenario. This aligns with our structural stability analysis, which identified the most stable configurations near average conditions. Third, an oscillating regime of alternating rainfall extremes also sustained biodiversity. As anticipated by our non-equilibrium framework, these continuous environmental fluctuations prevent the community from ever materializing the exclusions associated with a specific environment, effectively rescuing species through fluctuation-dependent mechanisms. In contrast, directional environmental changes led to community collapse. Rainfall sequences toward either increasing multi-year droughts (or positive anomalies) forced the system into persistently large fitness differences (or narrow feasibility domains), ultimately driving the simulations to competitive exclusion or total extinction.

The long-term simulations point the same way. That is, oscillating or constant rainfall scenarios provide the largest coexistence opportunities. They also show that context dependency (i.e., differences among sites) further modulates these outcomes. Since the mean environmental conditions of all scenarios was equal, the different outcomes of the simulations emerge from the differences in the covariance term, Eq. (7), that depends on the specific temporal sequences of environmental conditions and abundances. However, when we analyze the time-averaged systems, we do not find a one-to-one correspondence with the simulations. Some sites are predicted to have less feasible states than those recorded in simulations (Fig. S8, sites 2, 3, 4, 5, and 6), and some sites have more feasibility for rainfall scenarios with small feasibility domains (for example, increasing and decreasing rainfall in site 7). These results suggest that including the covariance of time-averaged systems, while capturing some of the temporal variability of precipitation and abundance, fails to describe the full spectrum of simulation results. Future work should discern which processes are not accounted for when using a covariance term in a time-average versus a time-varying framework.

That a time-averaged description falls short of the simulations is itself an argument for treating these communities as genuinely time-varying. Our approach conceptually aligns with fluctuation-dependent coexistence mechanisms, such as the storage effect, in which long-term persistence is achieved not by maintaining a static equilibrium but by adapting to the sequence of environmental conditions. More broadly, our work suggests that non-equilibrium dynamics can be theoretically approached by the study of instantaneous attractors. Although the formal application of this concept remains in its infancy in ecology (but see Chesson, 2017 for one single species), mathematical developments of non-autonomous dynamical systems provide a rigorous foundation for characterizing such instantaneous attractors (Carvalho et al., 2012; Bortolan et al., 2020; Godoy, Soler-Toscano, et al., 2024). These mathematical advances can provide a powerful predictive and explanatory framework for evaluating, for example, community responses to perturbations, designing targeted restoration trajectories, and anticipating the consequences of biological invasions. Unlike empirical forecasting methods based on machine learning or attractor reconstructions, e.g., S-maps Liu and Gaines, 2022; Ushio et al., 2018; Nguyen et al., 2025, that often preclude mechanistic interpretation and require fine-tuning, our tractable approach allows us to directly simulate population dynamics under hypothetical sequences of environmental change. More fundamentally, we advocate that capturing the complex dynamics of ecological communities will require, in many cases, embracing their inherently time-varying nature.

In sum, our study presents a readily available framework for obtaining a mechanistic understanding of the consequences of temporal environmental variation for species coexistence under natural conditions. Because the framework is general, it can be applied to any ecological system, other environmental drivers, and other sequences of time-varying interactions. Besides its theoretical advances, this framework opens the door to designing conservation strategies that help to forecast, and therefore, prevent biodiversity loss amid increased environmental anomalies.

## Acknowledgments

We thank Lauren Hallett for her insightful comments on the draft. We also thank Lisa Buche and María Hurtado de Mendoza for their help in collecting field data on annual plant abundances. OG and VC-S acknowledge support from the Spanish Ministry of Science and Innovation for the project PID2021-127607OB-I00. VC-S acknowledges Meteoblue for offering the historical rainfall data in the area (Aznalcazar station) free of charge.

