## Supplementary Information for "Time-varying interactions change coexistence mechanisms"

### Supplementary Information: Time-varying interactions change coexistence mechanisms

Violeta Calleja-Solanas[1,2], Ignasi Bartomeus[2] and Oscar Godoy[2]
[1]Department of Biology, University of Oxford, Oxford, United Kingdom
[2]Estación Biológica de Doñana (EBD-CSIC), Americo Vespucio 26 41092, Seville, Spain

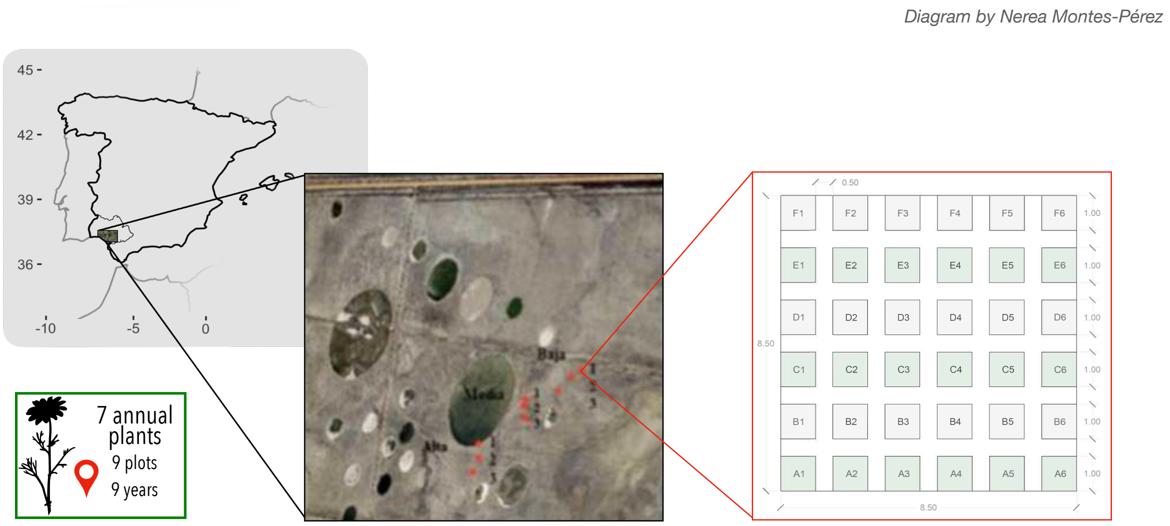


Figure 1: Satellite image of the 9 sampling sites, marked as red dots, and their design: Each plot is divided into subplots of 1 x 1 $m$. Within them, we measured the individuals of each focal species. We also left aisles of $0.5m$ between subplots for moving around without producing any disturbance. Figure provided by Nerea Montés Pérez. Grid inset modified from [1]

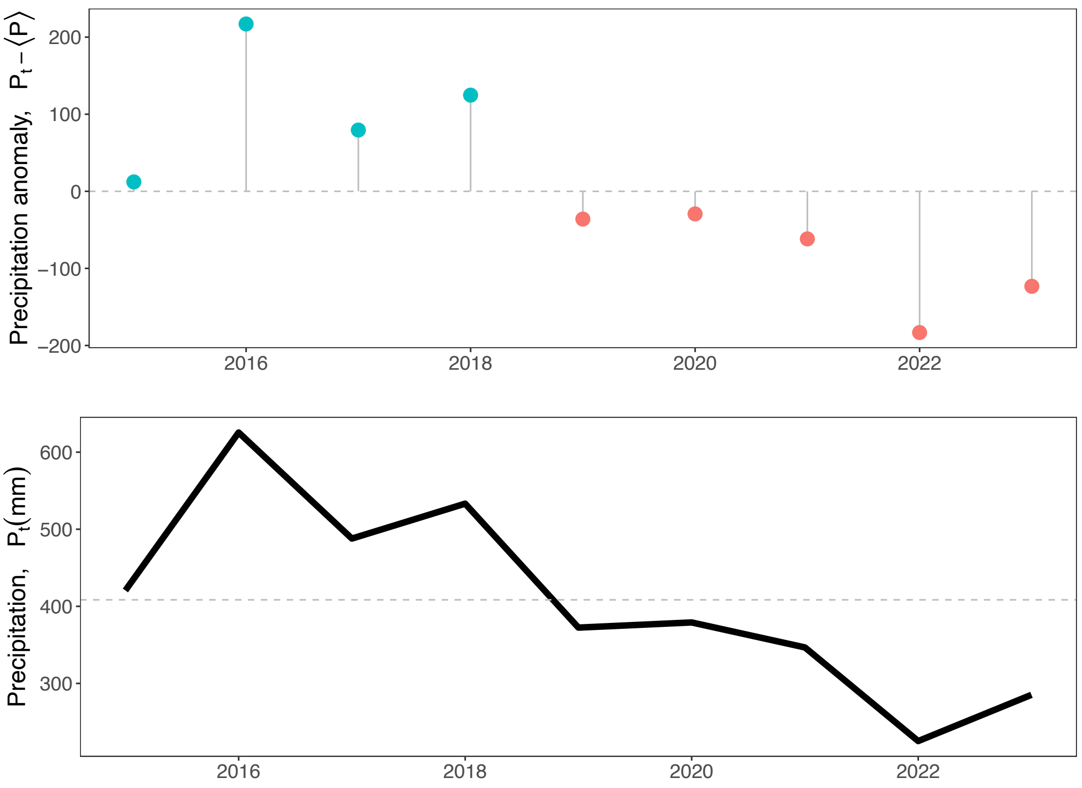


Figure 2: Total annual rainfall measured in Aznalcazar station, Spain, the nearest station to the sites. The 85-year rainfall average, $\langle\theta\rangle = 520\,mm$, was obtained from data of Meteoblue.

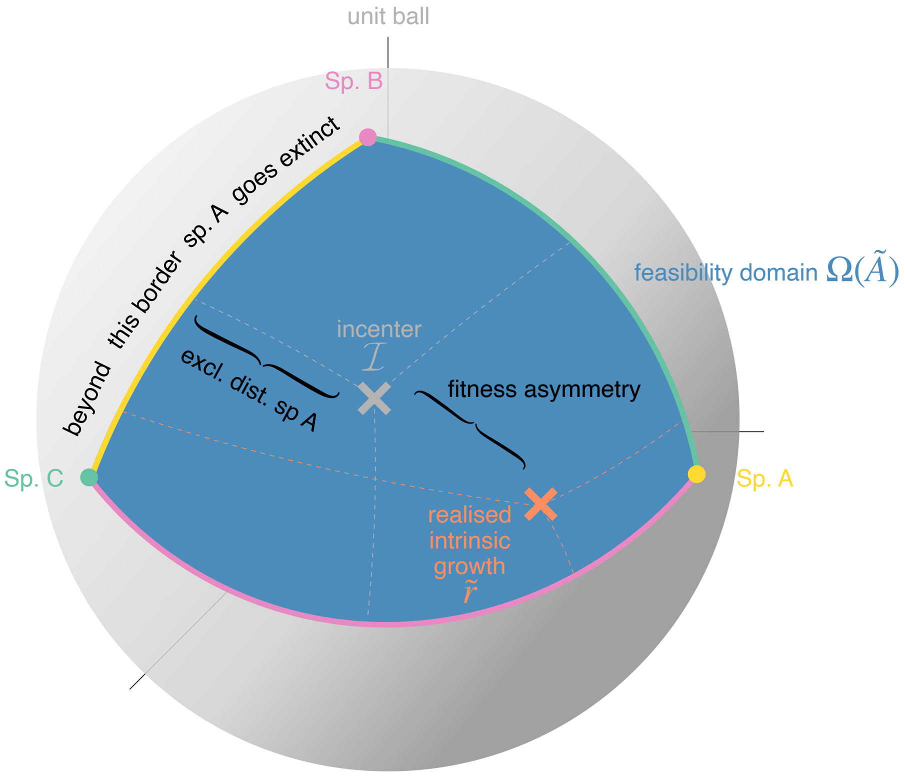


Figure 3: Diagram with the visual definitions of the measures of the structural stability approach for a 3-species system.

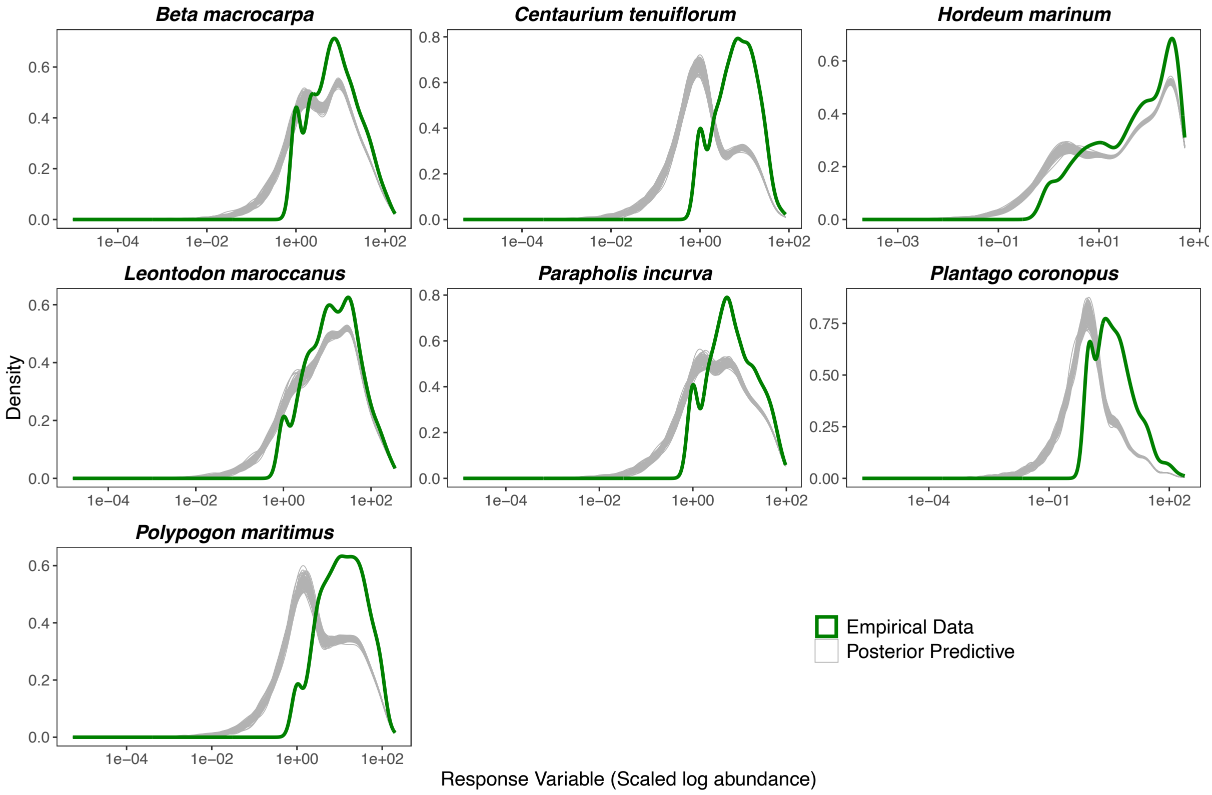


Figure 4: Posterior probability distribution of the normalized densities of species, obtained from the simulation of 1000 independent parameters draws.

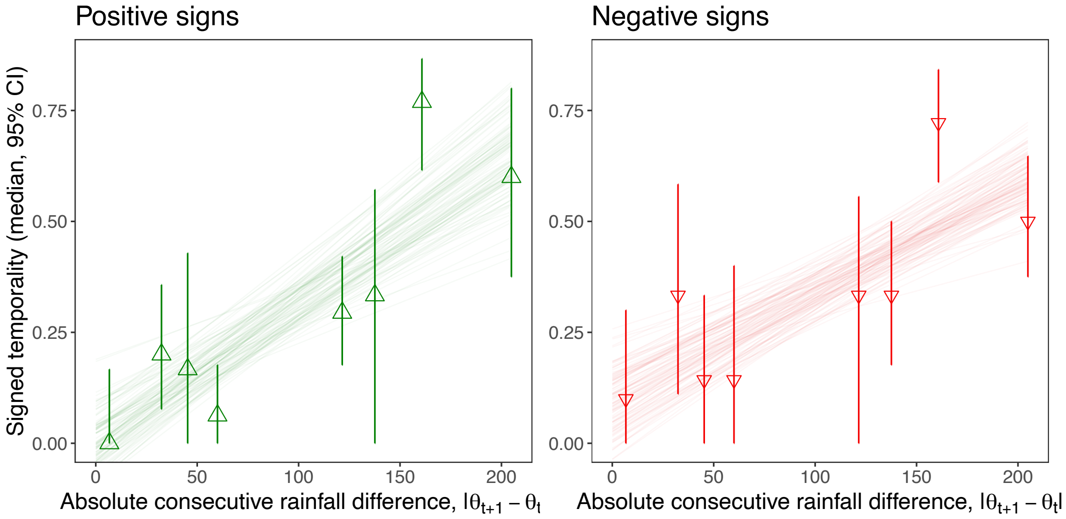


Figure 5: Variation in the temporality of only positive and only negative interactions: the extent to which two networks are similar over time, as a function of the absolute change in rainfall between two consecutive years, $|\theta_{t+1} - \theta_t|$. Points show the median across draws and vertical bars the $\pm 95\% CI$. The lines are least-squares fits of temporality against absolute rainfall difference obtained separately within 100 randomly chosen posterior draws, so that their spread reflects posterior uncertainty in the interaction parameters.

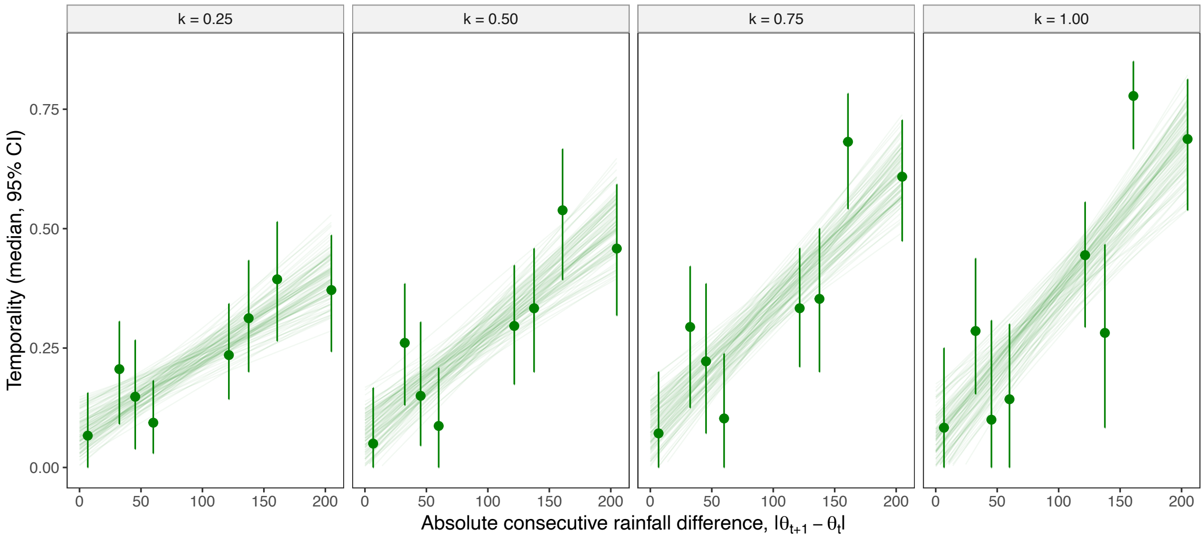


Figure 6: Temporality does not depend on the value of the scaling constant. Interaction coefficients were retained as edges when the standard deviation of the elements of the same matrix, and temporality was computed within each of the 1000 posterior draws. Posterior median and 95% credible interval against the absolute consecutive rainfall difference, for four values of $k$; lines are least-squares fits obtained separately within each of 100 randomly chosen draws.

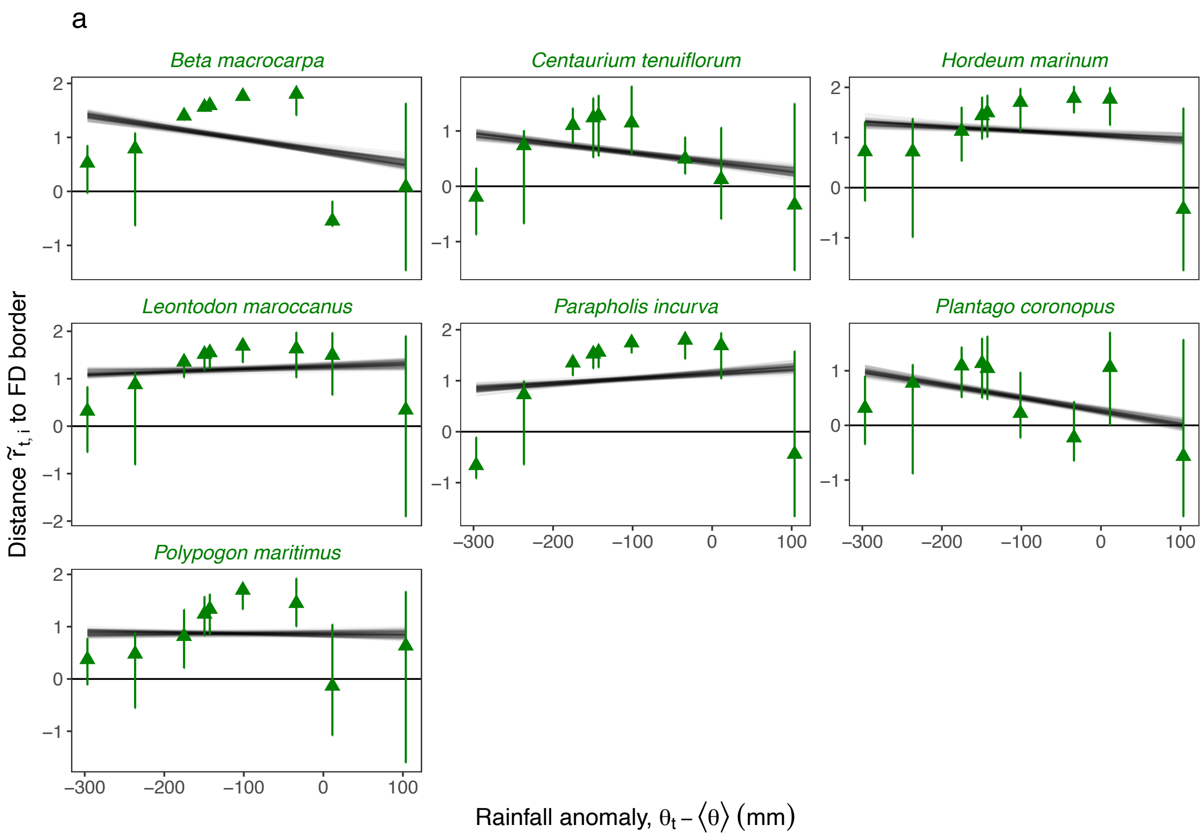


Figure 7: Distances of each species $i$ realized intrinsic growth rate $\tilde{r}_{t,b,i}$ to its extinction border. Each gray point is the median posterior distance from a particular draw of the species position in each site $b$ and time $t$. The slopes are the posterior mean estimates of the Bayesian linear regression of 1000 draws. They represent the overall tendency of a species reported as "Persistence" in the main text Fig. 4b.

Table 1: Species in the annual plant (Sites 1-9)

| **Species** |
| --- |
| *Beta macrocarpa* |
| *Centaurium tenuiflorum* |
| *Hordeum marinum* |
| *Leontodon maroccanus* |
| *Parapholis incurva* |
| *Polypogon maritimus* |

### 1 Mapping non-stationary dynamics to the framework of structural stability via time averaging

#### 1.1 Time-averaged system

We begin by recalling the dynamics equation:

$$\log \frac{N_{t+1}^{i,b}}{N_t^{i,b}} = r_i + r_i'\theta_t + u_{i,b} + \sum_{j=1}^{n}(A_{ij} + B_{ij}\theta_t)N_t^{j,b}. \tag{1}$$

Assuming the populations persist over a sufficiently long time, the abundance change (left-hand side of Eq. (1)) tends to zero after some transient:

$$\frac{1}{T}\sum_{t}^{T} \log \frac{N_{t+1}^{i,b}}{N_t^{i,b}} = \frac{\log N_{T+1}^{i,b} - \log N_1^{i,b}}{T} \approx 0 \tag{2}$$

Applying the time-averaging operator $\frac{1}{T}\sum_t^T$ to both sides, we denote the time average of a variable $x_t$ as $\overline{x}$. Averaging the right-hand side yields:

$$0 = r_i + r_i'\overline{\theta} + u_{i,b} + \sum_{j=1}^{n} A_{ij}\overline{N^{j,b}} + \sum_{j=1}^{n} B_{ij}\overline{\theta N^{j,b}}. \tag{3}$$

We expand the product term $\overline{\theta N^{j,b}}$ using $\overline{xy} = \overline{x}\cdot\overline{y} + \mathrm{cov}(x,y)$, obtaining:

$$0 = r_i + r_i'\overline{\theta} + u_{i,b} + \sum_{j=1}^{n}(A_{ij} + B_{ij}\overline{\theta})\overline{N^{j,b}} + \sum_{j=1}^{n} B_{ij}\mathrm{cov}(\theta, N^{j,b}). \tag{4}$$

We can map the latter expression into the classic linear form $\vec{r} = -\mathbf{A}\vec{N}^*$, by defining the effective time-averaged interaction matrix elements $\overline{\tilde{A}_{ij}}$ and the effective time-averaged intrinsic growth rates $\overline{\tilde{r}_i}$:

$$\overline{\tilde{A}_{ij}} = A_{ij} + B_{ij}\overline{\theta}, \tag{5}$$

$$\overline{\tilde{r}_i} = r_i + r_i'\overline{\theta} + u_{i,b} + \sum_{j=1}^{n} B_{ij}\mathrm{cov}(\theta, N^{j,b}). \tag{6}$$

Substituting these effective parameters back into the time-averaged equation gives the geometric decomposition for structural stability:

$$\overline{\tilde{r}_i} = -\sum_{j=1}^{n} \overline{\tilde{A}_{ij}}\,\overline{N^{j,b}}. \tag{7}$$

The covariance term acts as a environmental-driven fluctuation that can rotate the intrinsic growth rate vector across the feasibility domain.

#### 1.2 Consequences on structural stability

Since the effective time-averaged interaction matrix $\overline{\tilde{A}}$ only depends on the mean environmental conditions $\overline{\theta}$, two different sequences of environmental conditions will have the same feasibility domain (size and shape) if they share $\overline{\theta}$. On the contrary, time-averaged intrinsic growths $\overline{\tilde{r}_i}$ not only depend on $\overline{\theta}$, but also on a fluctuation-dependent term, the covariance between rainfall and the time series of species abundances. This covariance encodes the differences that arise from two different sequences of environmental conditions. Then, time-varying environmental conditions do not change coexistence opportunities, but can rotate the intrinsic growth rate vector across the feasibility domain via the covariance term.

To explore how different environmental conditions affect the structural stability of our community, we simulated different rainfall scenarios until they reach a stationary. Then, we calculated the time-averaged $\overline{\tilde{A}}$ and $\overline{\tilde{r}}$ (neglecting the transient), and compared their feasibility predictions with the simulations. In particular, we drawn 9000 parameters values from our posterior distributions and simulated five different rainfall scenarios: constant rainfall similar to the 85-year average, $\theta_{\mathrm{cte}} = 500$ mm, decreasing rainfall $\theta_{\downarrow}$ from 600 to 400 mm, increasing rainfall $\theta_{\uparrow}$ from 400 to 600 mm, a triangular wave $\theta_{\wedge}$ alternating between 600 and 400 mm, and a sinusoidal sequence $\theta_{\sim}$ centered at 500 mm with 100 mm of amplitude and a 5-year period.

The simulations with a $\theta_{\wedge}$ scenario obtained the most feasible communities at the end of our simulations (1762) than any other scenario, followed by $\theta_{\mathrm{cte}}$ (1534) and $\theta_{\sim}$ (1471). The decreasing and increasing scenarios had only 881 and 234 feasible communities, respectively. Interestingly, from these feasible communities, the time-averaged intrinsic growth rate overestimates and underestimates the feasibility of some scenarios depending on the sites. As a consequence, the time-averaged system does not capture the results of the simulation. The rainfall sequence is not fully captured in the covariance term.

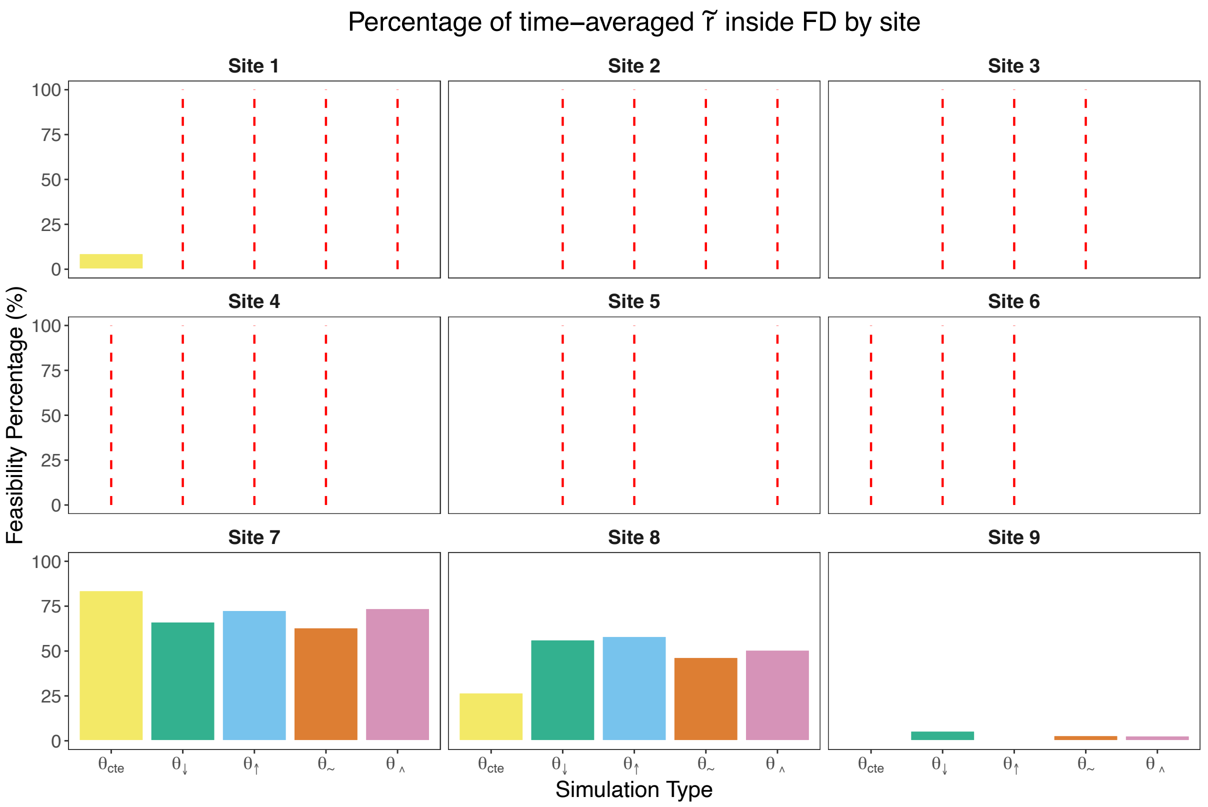


Figure 8: Percentage of time-averaged intrinsic growth rates $\bar{\tilde{r}}$ that were found inside the feasibility domain (FD) calculated from the time-average interactions $\bar{\tilde{A}}$. The simulation types are the different rainfall scenarios: constant rainfall $\theta_{\text{cte}} = 500$ mm, decreasing rainfall $\theta_{\downarrow}$ from 600 to 400 mm, increasing rainfall $\theta_{\uparrow}$ from 400 to 600 mm, a sinusoidal sequence $\theta_{\sim}$ centered at 500 mm with 100 mm of amplitude and a 5-year period, and a triangular wave $\theta_{\wedge}$ alternating between 600 and 400 mm. Red dashed lines indicate that no simulation ended up with full coexistence for a given site and rainfall scenario.

#### 2 Time-varying model

Among the different functional forms in which we can explicitly encode temporal variability on a parameter $p$ (a linear, nonlinear, sigmoid, exponential, or unimodal function), we chose a simple linear dependence of the form $\tilde{p}(t) = mf(t) + p_0$. That is, the now time-varying parameter grows from a baseline $p_0$ linearly. The time-dependent function $f(t)$ encodes an environmental driver that varies across time, such as rainfall or temperature, and $m$ is the effect of its value on the parameter. For our study systems, $m = r_i'$ and $B_{ij}$ for intrinsic growth rates and interactions, respectively, and $f(t) = \theta_t$, the annual rainfall at time $t$ (Fig. 2). We can interpret this linear dependency as a first-order Taylor expansion of any other functional form. Accordingly, the model is valid only in a regime where the environmental variation is not extreme. Moreover, there exists a threshold value $p_c$ for which $\tilde{p}(t)$ intersects the abscissa. Crucially, if that value is within the typical environmental variation, then our parameter will change sign (Fig. 9).

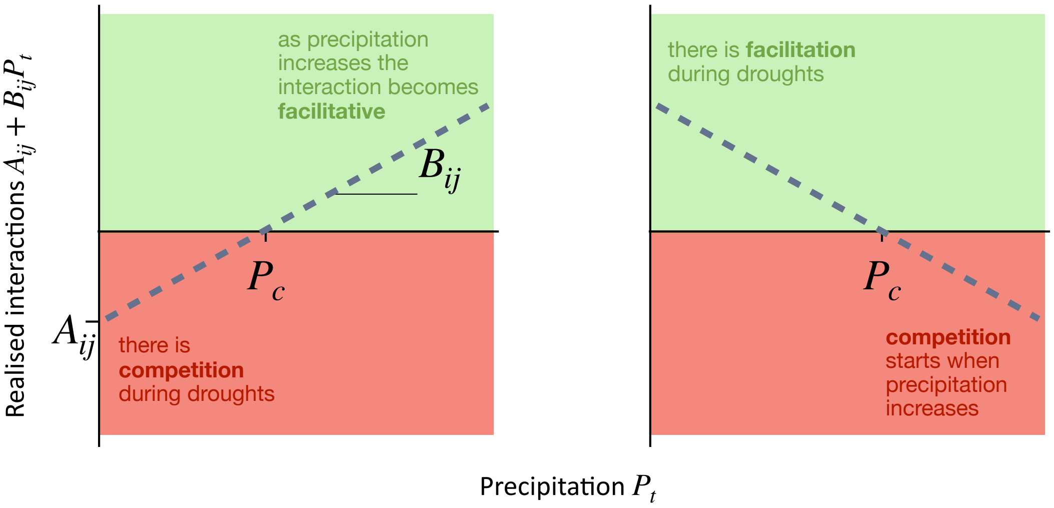


Figure 9: Example of time-varying precipitation $P_t$ behavior.

We found that the threshold environmental values ($pc$) at which intrinsic growth rates and species interactions change sign are largely contained within the range of typical historical environmental variation (Fig. 10). To compute $pc$, we used the means of the posterior distributions. Specifically, $pc$ represents the critical rainfall value $\theta_t$ at which a net interaction ($\tilde{A}_{ij} = A_{ij} + B_{ij}\theta_t$) or an intrinsic growth rate ($\tilde{r}_i = r_i + r_i'\theta_t$) crosses zero, shifting between positive and negative effects.

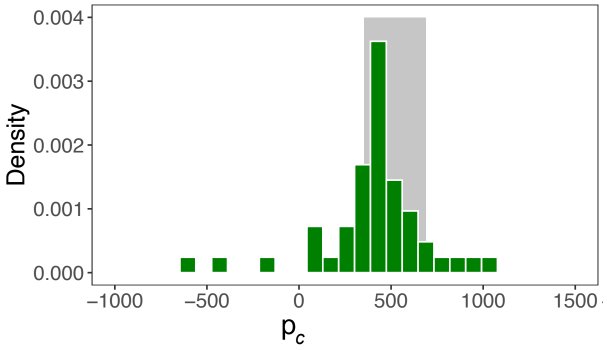


Figure 10: Histogram of the sign-change thresholds ($p_c$) for each interaction element and intrinsic growth rate. The gray ribbon indicates the historical rainfall variation over the last 85 years (spanning one standard deviation from the mean).

#### 2.1 Bayesian model selection

To find a compromise between the number of model parameters (which limits computational tractability and requires rich longitudinal data) and the mechanistic representation of temporal variability, we formalized a set of competing models.

The selected model (Eq. 2 in the main text) incorporates temporal variability in both interactions and intrinsic growth rates coefficients:

$$\log \frac{N_{t+1}^{i,b}}{N_t^{i,b}} = r_i + r_i'\theta_t + \sum_{j=1}^{n}(A_{ij} + B_{ij}\theta_t)N_t^{j,b} + u_{i,b}, \tag{8}$$

where $u_{i,b}$ is the hierarchical, site-level stochastic variation.

We compared this full framework against simplified models. For example,

a model of only time-varying interactions is formulated as:

$$\log \frac{N_{t+1}^{i,b}}{N_t^{i,b}} = r_i + \sum_{j=1}^{n}(A_{ij} + B_{ij}\theta_t)N_t^{j,b} + u_{i,b}. \tag{9}$$

Alternatively, a model where temporal variability is restricted to intrinsic growth rates:

$$\log \frac{N_{t+1}^{i,b}}{N_t^{i,b}} = r_i + r_i'\theta_t + \sum_{j=1}^{n} A_{ij}N_t^{j,b} + u_{i,b} + u_{i,b}'\theta_t, \tag{10}$$

in which there is also a stochastic effect on the rainfall response for the intrinsic growth rate $u_{i,b}'$. Note that we denote the effects of temporal variability on different aspects of intrinsic growth with prime notation.

To evaluate these competing models, we calculated the out-of-sample predictive performance using approximate leave-one-out cross-validation (LOO-CV) via the R package `loo`, Table 3.

##### 2.1.1 Bayesian model setting

To estimate the posterior distributions of the parameters $\{r_i, r_i', A_{ij}, B_{ij}\}$, we fitted the models using Hamiltonian Monte Carlo (HMC) sampling via the R package `brms` (with `cmdstanr` backend). We modeled the log-transformed population growth rates using a Gaussian likelihood. To prevent overfitting and ensure robust parameter estimation, we applied weakly informative regularizing priors: specifically, we assigned a Normal prior $\mathcal{N}(0, 0.5)$ to all population-level effects, and a half-Normal prior $\mathcal{H}(0, 1)$ to the standard deviations of the hierarchical site-level effects, $u_{i,b}$. Models were run across four parallel chains with 2000 iterations each (plus 1000 warm-up iterations), obtaining a total of 4000 posterior samples. Because our time-varying and hierarchical structure creates a complex posterior geometry, we increased the `adapt_delta` parameter to 0.99 to eliminate divergent transitions, and raised the `max_treedepth` to 12 to ensure efficient exploration of the parameter space.

Table 2: Description of Leave-One-Out Cross-Validation (LOO-CV) metrics for Bayesian model comparison.

| Metric | Description |
|---|---|
| `elpd_loo` | **Expected Log Predictive Density:** The estimate of the model's out-of-sample predictive accuracy. Higher (less negative) values indicate a better-fitting model with higher predictive power. |
| `se_elpd_loo` | The standard error of the expected log predictive density. |
| `p_loo` | **Effective Number of Parameters:** A measure of the model's estimated complexity. It acts as a penalization term against overfitting. |
| `se_p_loo` | The standard error of the effective number of parameters. |
| `looic` | **Leave-One-Out Information Criterion:** Calculated as $-2\times$elpd_loo. This metric is on the same deviance scale as AIC or WAIC. Lower values indicate better predictive performance. |
| `se_looic` | The standard error of the LOOIC. |
| `elpd_diff` | **$\Delta$ELPD:** The difference in expected log predictive density between a given model and the best-fitting model in the candidate set. By definition, the best model has an `elpd_diff` of 0. |
| `se_diff` | **Standard error of component-wise differences:** The standard error of the difference in predictive performance (`elpd_diff`). Because models are evaluated on the exact same dataset, their pointwise predictive densities are correlated; this metric accounts for that correlation. |
| `signif_worse` | A model is generally considered significantly worse than the best model if its absolute `elpd_diff` is more than twice its `se_diff`. Conversely, models are considered equivalent if they rest within that range. |

Table 3: Model comparison results for the seven species using Leave-One-Out Cross-Validation (LOO-CV). Column names' abbreviations are defined in Table 2. Models' abbreviations: Full = Eq. (8) and Eq. 2 in main text, Time-var A = Eq. (9), Time-var r = Eq. (10), Static = Eq. 1 in main text.

| ***Sp***, Model | `elpd_diff` | `se_diff` | `elpd_loo` | `se_elpd_loo` | `p_loo` | `se_p_loo` | `looic` | `se_looic` | `signif_worse` |
|---|---|---|---|---|---|---|---|---|---|
| ***Beta macrocarpa*** | | | | | | | | | |
| Full | 0.00 | 0.00 | -3495.32 | 38.80 | 27.35 | 1.25 | 6990.64 | 77.60 | F |
| Time-var A | -176.95 | 16.29 | -3672.27 | 36.60 | 20.42 | 1.03 | 7344.55 | 73.20 | T |
| Time-var r | -286.08 | 23.98 | -3781.40 | 40.10 | 18.18 | 1.14 | 7562.80 | 80.20 | T |
| Static | -839.86 | 30.79 | -4335.18 | 29.57 | 14.42 | 0.45 | 8670.35 | 59.14 | T |
| ***Centaurium tenuiflorum*** | | | | | | | | | |
| Full | 0.00 | 0.00 | -3309.51 | 55.16 | 43.60 | 5.73 | 6619.02 | 110.32 | F |
| Time-var A | -72.35 | 12.88 | -3381.87 | 56.43 | 35.50 | 5.23 | 6763.73 | 112.86 | T |
| Time-var r | -147.90 | 18.81 | -3457.41 | 53.73 | 24.69 | 3.06 | 6914.82 | 107.46 | T |
| Static | -190.67 | 23.13 | -3500.18 | 56.43 | 23.49 | 3.04 | 7000.36 | 112.87 | T |
| ***Hordeum marinum*** | | | | | | | | | |
| Full | 0.00 | 0.00 | -4771.49 | 40.84 | 26.10 | 0.99 | 9542.97 | 81.67 | F |
| Time-var A | -32.49 | 6.89 | -4803.97 | 41.18 | 20.12 | 0.80 | 9607.95 | 82.35 | T |
| Time-var r | -243.26 | 18.36 | -5014.74 | 36.68 | 16.87 | 0.53 | 10029.48 | 73.36 | T |
| Static | -407.26 | 21.66 | -5178.74 | 37.75 | 15.67 | 0.50 | 10357.48 | 75.49 | T |
| ***Leontodon maroccanus*** | | | | | | | | | |
| Full | 0.00 | 0.00 | -4362.51 | 35.99 | 32.79 | 1.47 | 8725.03 | 71.97 | F |
| Time-var A | -117.64 | 16.01 | -4480.16 | 35.61 | 23.97 | 1.16 | 8960.32 | 71.22 | T |
| Time-var r | -309.41 | 20.74 | -4671.92 | 35.89 | 19.20 | 1.04 | 9343.85 | 71.78 | T |
| Static | -450.54 | 25.87 | -4813.05 | 38.06 | 17.93 | 1.05 | 9626.10 | 76.12 | T |
| ***Parapholis incurva*** | | | | | | | | | |
| Full | 0.00 | 0.00 | -4060.52 | 36.22 | 36.06 | 2.72 | 8121.04 | 72.44 | F |
| Time-var A | -46.54 | 11.40 | -4107.06 | 34.87 | 28.05 | 1.96 | 8214.12 | 69.74 | T |
| Time-var r | -331.57 | 23.80 | -4392.09 | 32.71 | 20.03 | 1.08 | 8784.18 | 65.42 | T |
| Static | -452.34 | 27.40 | -4512.86 | 32.93 | 18.82 | 1.03 | 9025.73 | 65.85 | T |
| ***Plantago coronopus*** | | | | | | | | | |
| Full | 0.00 | 0.00 | -3304.63 | 64.59 | 46.31 | 7.46 | 6609.26 | 129.17 | F |
| Time-var A | -22.85 | 7.98 | -3327.48 | 64.54 | 39.60 | 7.08 | 6654.96 | 129.08 | T |
| Time-var r | -155.88 | 20.77 | -3460.51 | 65.70 | 33.52 | 6.66 | 6921.01 | 131.41 | T |
| Static | -157.30 | 21.38 | -3461.93 | 65.83 | 32.78 | 6.89 | 6923.85 | 131.65 | T |
| ***Polypogon maritimus*** | | | | | | | | | |
| Full | 0.00 | 0.00 | -4303.44 | 40.10 | 35.86 | 2.67 | 8606.89 | 80.20 | F |
| Time-var A | -206.39 | 20.22 | -4509.83 | 37.74 | 24.27 | 2.10 | 9019.66 | 75.49 | T |
| Time-var r | -371.82 | 31.83 | -4675.26 | 39.07 | 19.54 | 1.57 | 9350.53 | 78.15 | T |
| Static | -452.46 | 30.13 | -4755.91 | 36.79 | 17.33 | 1.46 | 9511.82 | 73.58 | T |

### 3 Time-varying framework with GLME

To verify that our ecological conclusions are not artifacts of the chosen Bayesian priors or inferring method, we additionally fitted the time-varying framework using a frequentist approach: generalized linear mixed-effects models (GLME) via the R package `nlme`. We modeled the log-transformed population growth rates using Restricted Maximum Likelihood (REML), which provides unbiased estimates of variance and covariance parameters. Mirroring our Bayesian structure, the models incorporated fixed effects for baseline intrinsic growth rates, species interactions, and their continuous responses to rainfall. To account for local spatial contingencies, we included hierarchical random intercepts for the sampling sites and random slopes for the rainfall effect at each site.

#### 3.1 Results comparison between GLME and Bayesian inference

This dual-inference approach ensures that our estimates of time-varying interactions and intrinsic growth rates are structurally robust, since we found the same patterns of change and reorganization. In particular, the estimated intrinsic growth rates and interactions coincide with the median of the posterior distributions (Fig. 11). This generates the same patterns of change we found on the main text for the temporality (Fig. 12), interspecific interactions and diagonal dominance (Fig. 13). The trends also combine to increase coexistence opportunities (Fig. 12) and fitness differences (Fig. 14) during negative rainfall anomalies (Fig. 15). However, despite these overarching similarities, the nonlinear shifts in structural stability captured by the Bayesian model were notably less pronounced under the GLME framework (Fig. 16).

All estimated parameters are provided in Supplementary Information 3.2, and the validation of predicted abundances can be found in Fig. 17.

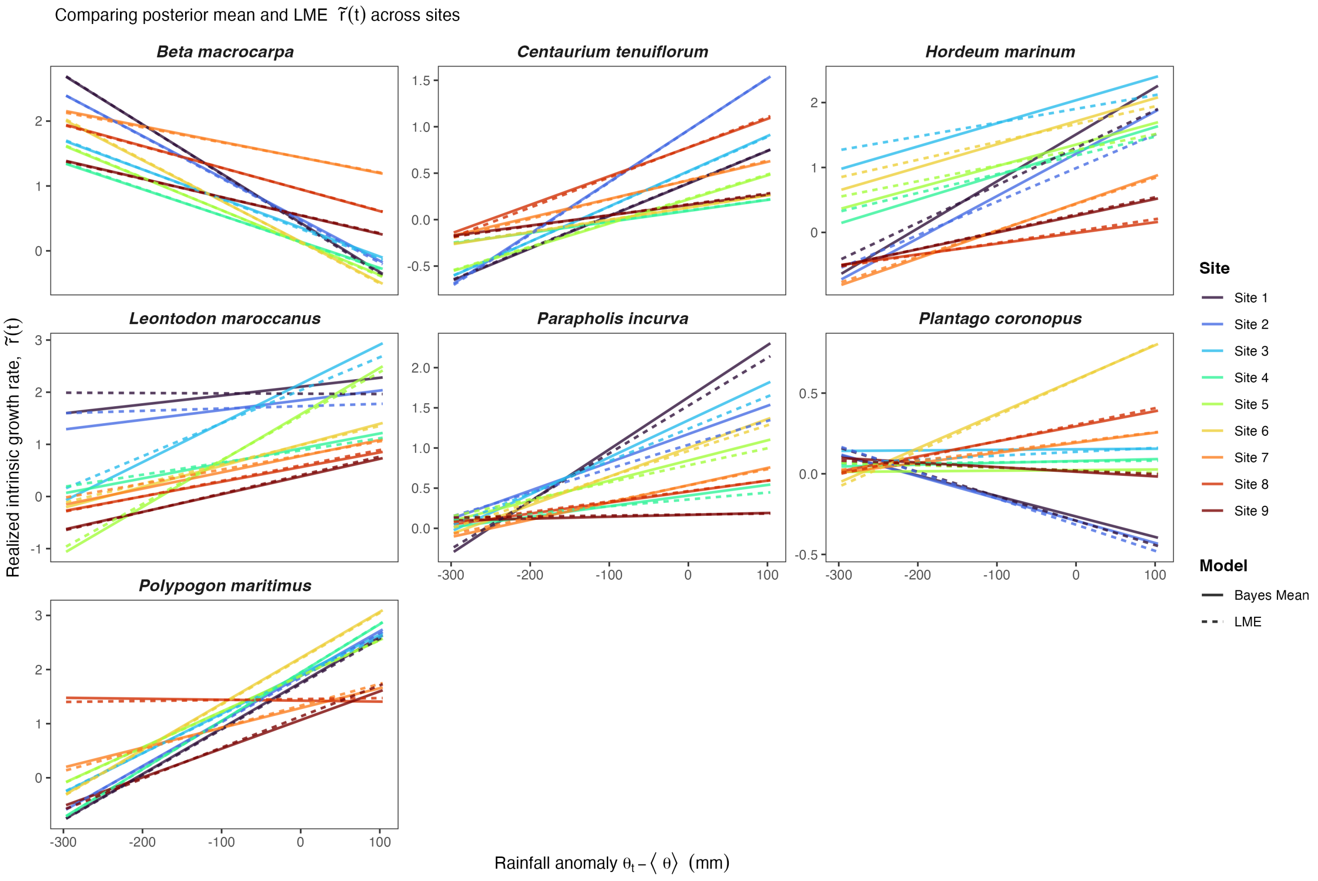


Figure 11: Comparison between the realized intrinsic growth rate $\tilde{r}(t)$ inferred with the GLME and the mean of the posterior distribution obtained by Bayesian inference.

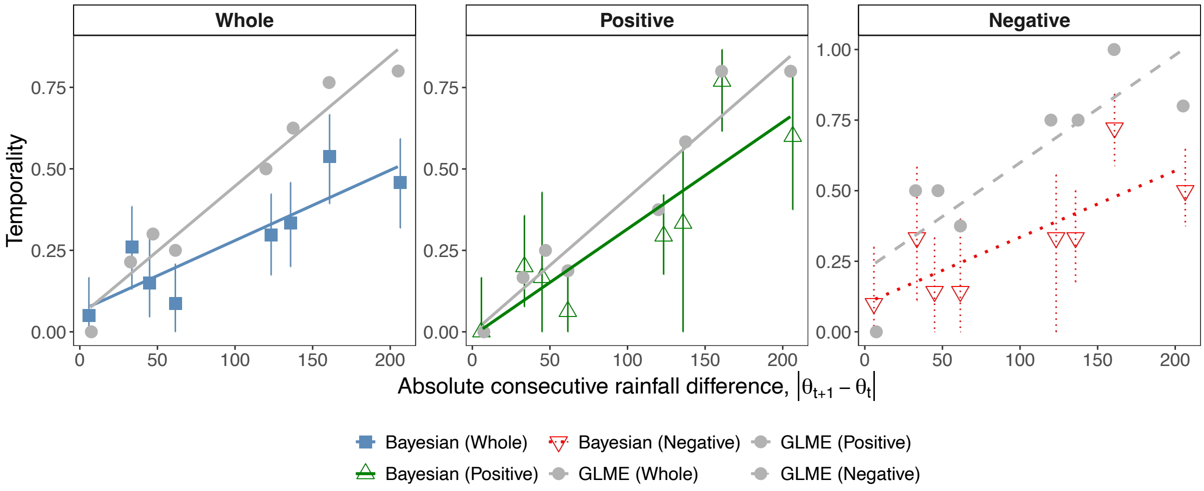


Figure 12: Comparison between the signed temporality of the networks inferred with the GLME method and Bayesian inference. Jitter was added to the points' x-coordinates to improve visualization. Lines are the least-squares regression trends of the points (not jittered).

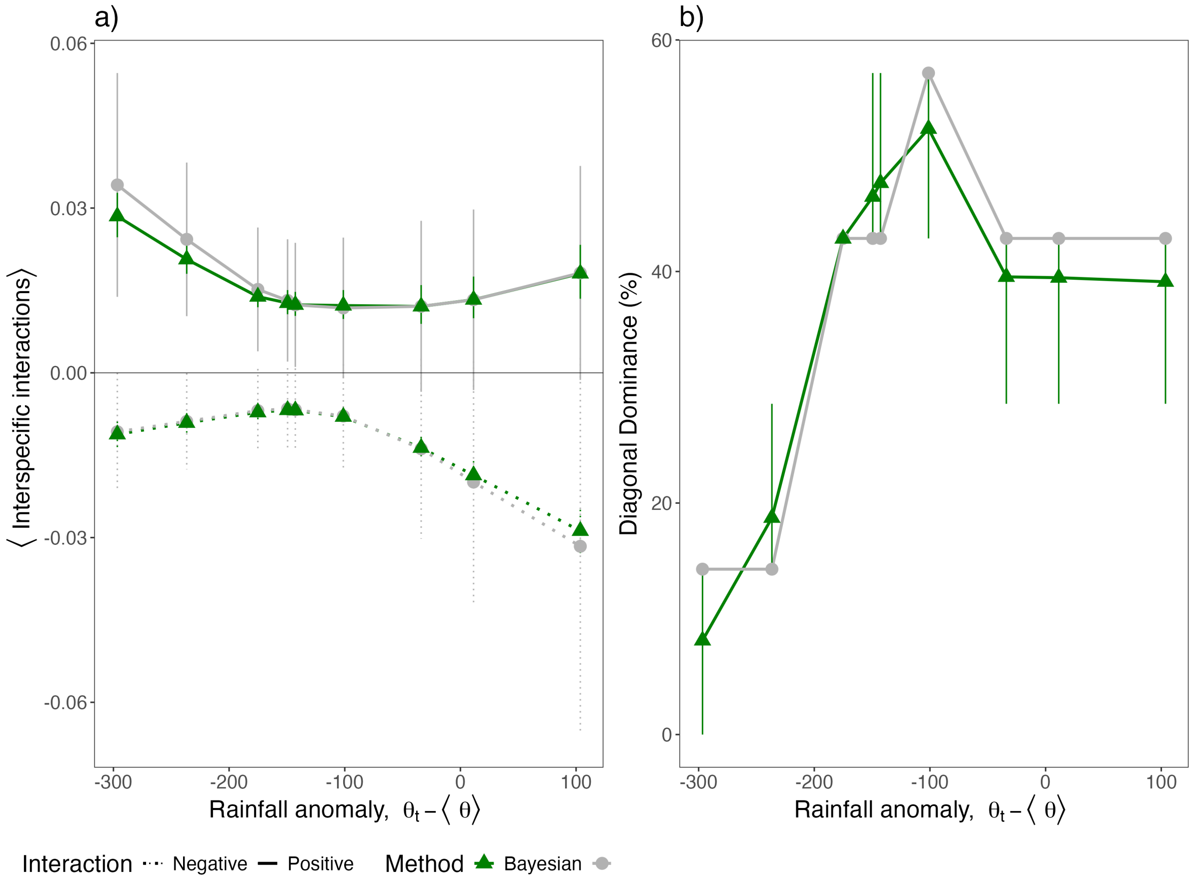

a)
b)
⟨ Interspecific interactions⟩
Diagonal Dominance (%)
Rainfall anomaly, θt − ⟨ θ⟩
Rainfall anomaly, θt − ⟨ θ⟩
Interaction Negative Positive Method Bayesian


Figure 13: (a) (b)

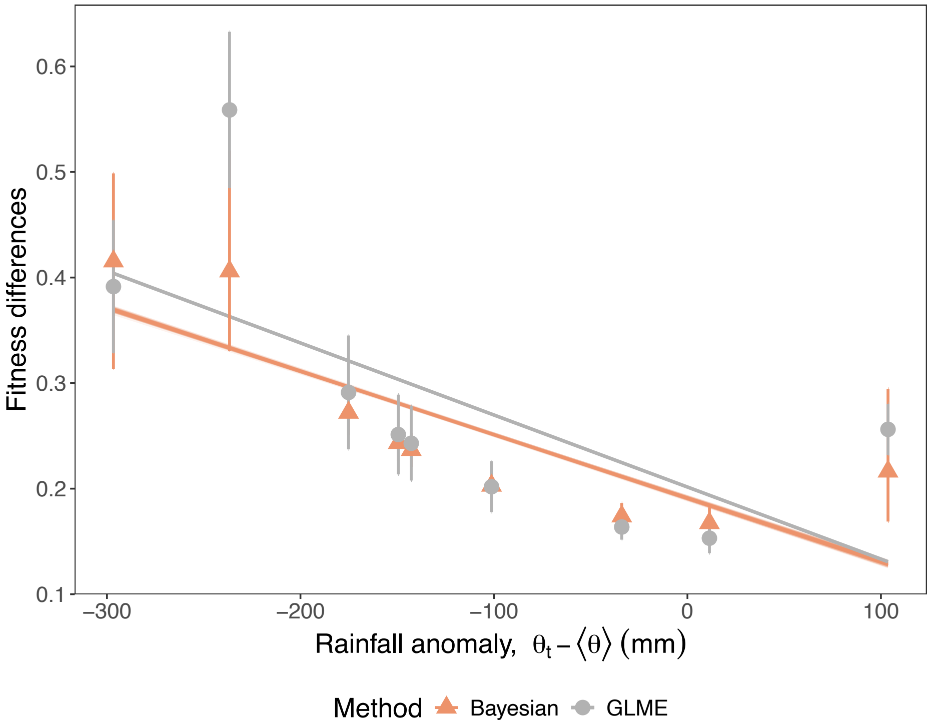


Figure 14: Estimated fitness differences within the community and sites along the rainfall gradient. Orange triangles represent Bayesian posterior means with 95% credible intervals, overlaid on faint blue lines representing trends of individual posterior draws. Gray circles indicate GLME estimates, with error bars representing ±1 standard deviation, and the solid gray line illustrating the least-squares linear trend.

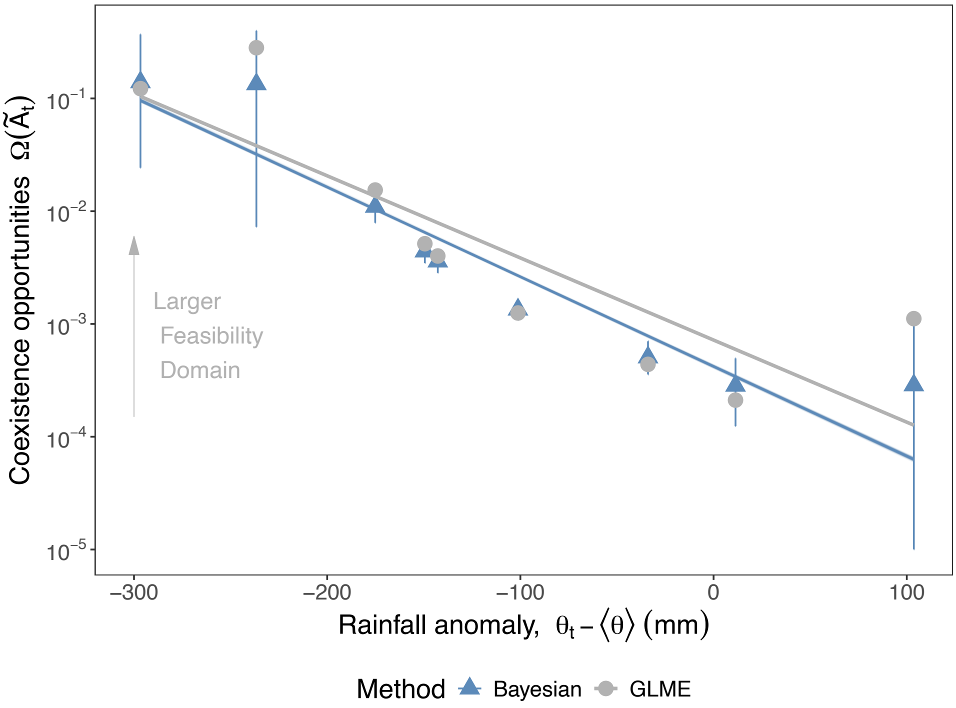


Figure 15: Comparison of estimated coexistence opportunities across a rainfall gradient. The size of the feasibility domain, $\Omega(\tilde{A}(t)$, represents the probability of coexistence under given interaction constraints. Blue triangles denote Bayesian posterior means with 95% Credible Intervals. The faint background lines (which are overlapping) represent posterior draws to illustrate estimation uncertainty. Gray circles denote Generalized Linear Mixed-Effects (GLME) model estimates, with the solid gray line representing the least-squares linear trend.

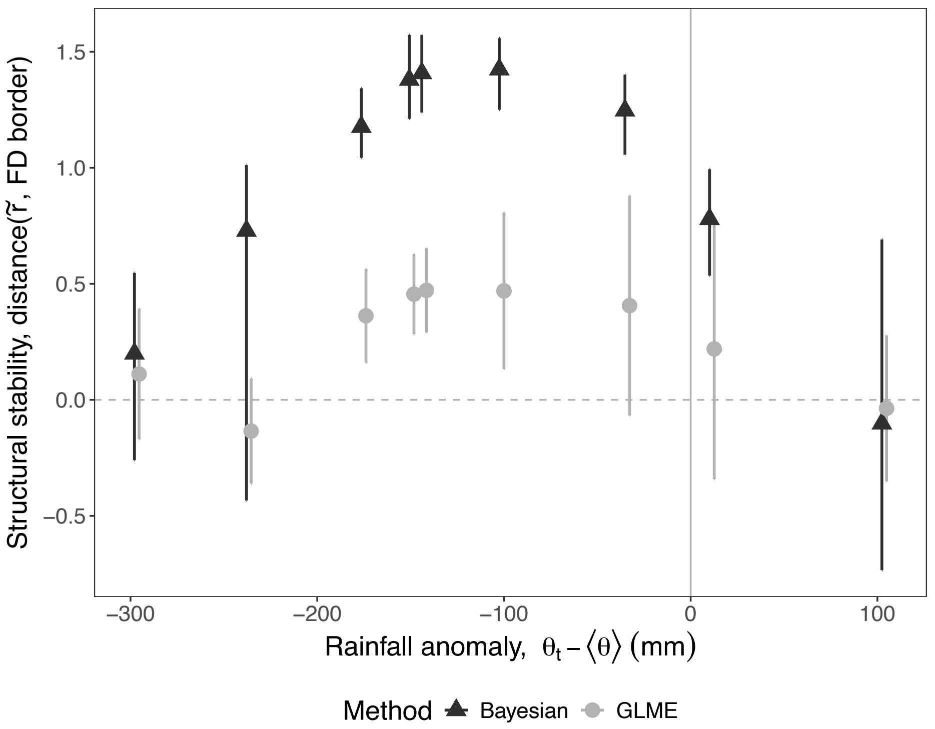


Figure 16: Structural stability is quantified as the distance of the intrinsic growth rate vector ($\tilde{r}$) to the borders of the feasibility domain. Triangles (Bayesian posterior means ±95% CI) and gray circles (GLME estimates ±1 standard deviation, scaled by $\pi$) represent the respective models estimates. Points are horizontally jittered to prevent overlapping error bars. The horizontal dashed line at zero indicates the threshold where some species is out of the feasibility domain.

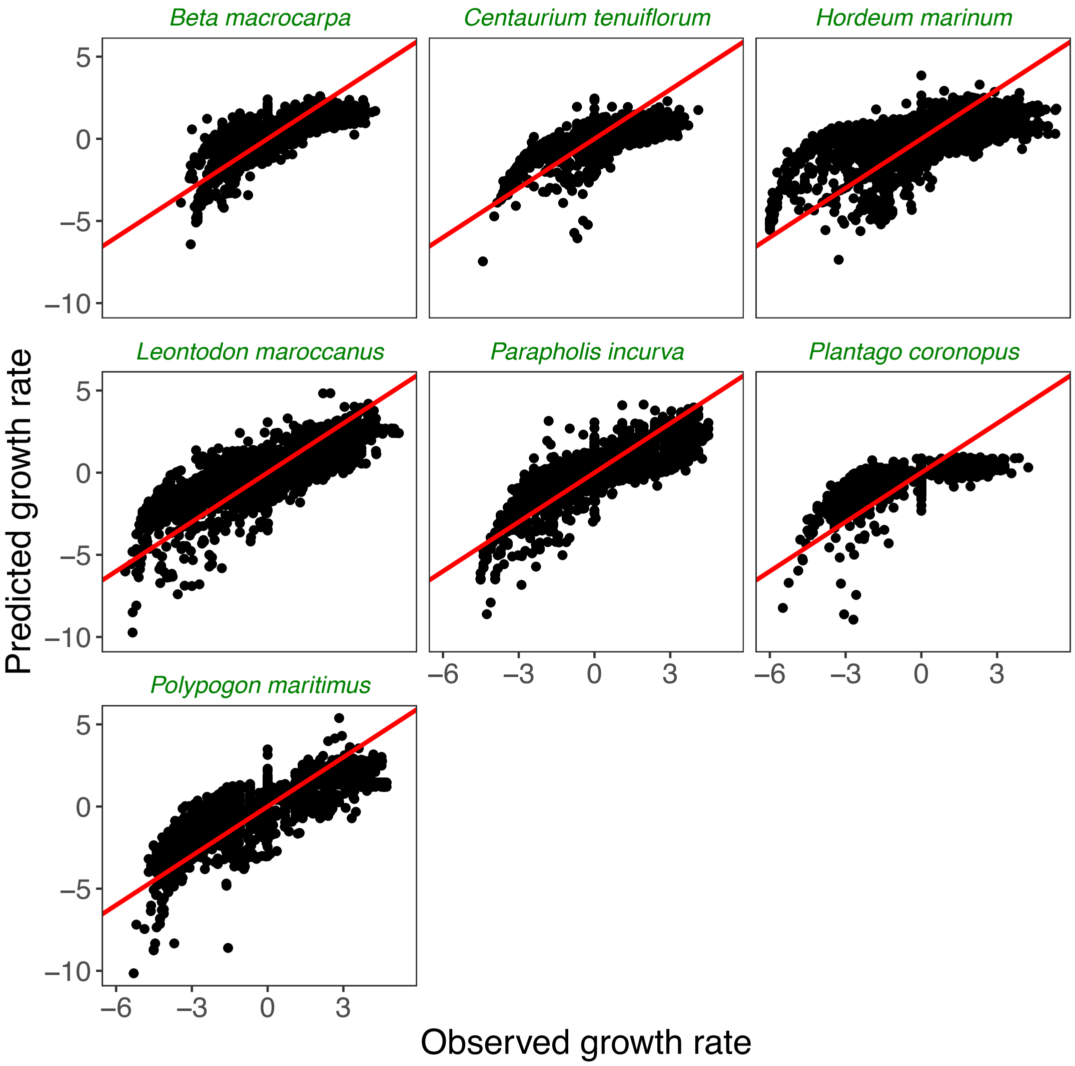


Figure 17: Scatter plots of the observed and predicted growth rates obtained by integrating Eq. (8) with the parameter values of each time $t$ and site $b$. Red lines have slope one. Observed values were computed from species overall abundance (individuals counted), and predictions are simulated values with the empirical rainfalls.

#### 3.2 Estimated parameters for the GLME model

Ready-to-use files with the values are in Zenodo repository https://zenodo.org/records/15625585.

##### 3.2.1 Realized interaction matrix $\tilde{A}_t = A + B\theta_t$

$$A = \begin{pmatrix} 0.0774 & 0.0201 & -0.0016 & -0.0011 & 0.1095 & -0.0206 & -0.0008 \\ 0.0397 & -0.0531 & 0.0055 & -0.0201 & 0.0739 & -0.0421 & 0.0939 \\ 0.0300 & -0.0747 & -0.0400 & -0.0319 & -0.0076 & 0.0160 & 0.1006 \\ 0.0645 & 0.0656 & -0.0110 & -0.1195 & 0.0978 & 0.0073 & 0.1887 \\ 0.0228 & 0.0726 & -0.0270 & -0.0027 & 0.1867 & -0.0376 & 0.1057 \\ -0.0266 & 0.0601 & -0.0050 & -0.0003 & -0.0147 & -0.0899 & 0.1276 \\ 0.1677 & 0.0064 & -0.0011 & -0.0250 & 0.0555 & -0.0552 & 0.1668 \end{pmatrix} \quad \begin{matrix} \textit{Beta macrocarpa} \\ \textit{Centaurium tenuiflorum} \\ \textit{Hordeum marinum} \\ \textit{Leontodon maroccanus} \\ \textit{Parapholis incurva} \\ \textit{Plantago coronopus} \\ \textit{Polypogon maritimus} \end{matrix}$$

$$sd(A) = \begin{pmatrix} 0.00003 & 0.00005 & 0.00000 & 0.00001 & 0.00005 & 0.00003 & 0.00002 \\ 0.00003 & 0.00005 & 0.00000 & 0.00001 & 0.00004 & 0.00002 & 0.00002 \\ 0.00005 & 0.00009 & 0.00000 & 0.00001 & 0.00007 & 0.00004 & 0.00004 \\ 0.00004 & 0.00007 & 0.00000 & 0.00001 & 0.00006 & 0.00003 & 0.00003 \\ 0.00004 & 0.00006 & 0.00000 & 0.00001 & 0.00006 & 0.00003 & 0.00003 \\ 0.00003 & 0.00005 & 0.00000 & 0.00001 & 0.00004 & 0.00002 & 0.00002 \\ 0.00004 & 0.00007 & 0.00000 & 0.00001 & 0.00006 & 0.00003 & 0.00003 \end{pmatrix} \quad \begin{matrix} \textit{Beta macrocarpa} \\ \textit{Centaurium tenuiflorum} \\ \textit{Hordeum marinum} \\ \textit{Leontodon maroccanus} \\ \textit{Parapholis incurva} \\ \textit{Plantago coronopus} \\ \textit{Polypogon maritimus} \end{matrix}$$

$$B = \begin{pmatrix} -0.00066 & -0.00007 & 0.00000 & 0.00000 & -0.00025 & 0.00004 & 0.00000 \\ -0.00008 & -0.00011 & -0.00001 & 0.00005 & -0.00016 & 0.00009 & -0.00022 \\ -0.00007 & 0.00019 & 0.00006 & 0.00006 & 0.00011 & -0.00003 & -0.00022 \\ -0.00010 & -0.00017 & 0.00002 & 0.00020 & -0.00020 & -0.00004 & -0.00047 \\ 0.00004 & -0.00021 & 0.00006 & -0.00000 & -0.00066 & 0.00005 & -0.00028 \\ 0.00008 & -0.00014 & 0.00001 & 0.00000 & 0.00006 & 0.00011 & -0.00033 \\ -0.00049 & -0.00009 & -0.00001 & 0.00005 & -0.00015 & 0.00008 & -0.00055 \end{pmatrix} \quad \begin{matrix} \textit{Beta macrocarpa} \\ \textit{Centaurium tenuiflorum} \\ \textit{Hordeum marinum} \\ \textit{Leontodon maroccanus} \\ \textit{Parapholis incurva} \\ \textit{Plantago coronopus} \\ \textit{Polypogon maritimus} \end{matrix}$$

$$sd(B) = \begin{pmatrix} 0.00003 & 0.00005 & 0.00000 & 0.00001 & 0.00005 & 0.00003 & 0.00002 \\ 0.00003 & 0.00005 & 0.00000 & 0.00001 & 0.00004 & 0.00002 & 0.00002 \\ 0.00005 & 0.00009 & 0.00000 & 0.00001 & 0.00007 & 0.00004 & 0.00004 \\ 0.00004 & 0.00007 & 0.00000 & 0.00001 & 0.00006 & 0.00003 & 0.00003 \\ 0.00004 & 0.00006 & 0.00000 & 0.00001 & 0.00006 & 0.00003 & 0.00003 \\ 0.00003 & 0.00005 & 0.00000 & 0.00001 & 0.00004 & 0.00002 & 0.00002 \\ 0.00004 & 0.00007 & 0.00000 & 0.00001 & 0.00006 & 0.00003 & 0.00003 \end{pmatrix} \quad \begin{matrix} \textit{Beta macrocarpa} \\ \textit{Centaurium tenuiflorum} \\ \textit{Hordeum marinum} \\ \textit{Leontodon maroccanus} \\ \textit{Parapholis incurva} \\ \textit{Plantago coronopus} \\ \textit{Polypogon maritimus} \end{matrix}$$

##### 3.2.2 Realized intrinsic growth rates

Note that the values in $r'_i$ will be multiplied by rainfall $\theta_t \in (100, 700)$ mm to obtain $\tilde{r}_{t,b,i}$, Eq. (??).

Table 4: Site-level random intercepts and rainfall effects for each species.

| **Species** | **Site** | $r_i$ | $r'_i$ | **Std.Error** $r_i$ | **Std.Error** $r'_i$ | $u_{i,b}$ |
|---|---|---|---|---|---|---|
| *Beta macrocarpa* | 1 | 2.98 | -0.00474 | 0.296 | 0.000689 | 1.44 |
| *Beta macrocarpa* | 2 | 2.98 | -0.00474 | 0.296 | 0.000689 | 0.874 |
| *Beta macrocarpa* | 3 | 2.98 | -0.00474 | 0.296 | 0.000689 | -0.247 |
| *Beta macrocarpa* | 4 | 2.98 | -0.00474 | 0.296 | 0.000689 | -0.704 |
| *Beta macrocarpa* | 5 | 2.98 | -0.00474 | 0.296 | 0.000689 | -0.224 |
| *Beta macrocarpa* | 6 | 2.98 | -0.00474 | 0.296 | 0.000689 | 0.477 |
| *Beta macrocarpa* | 7 | 2.98 | -0.00474 | 0.296 | 0.000689 | -0.329 |
| *Beta macrocarpa* | 8 | 2.98 | -0.00474 | 0.296 | 0.000689 | -0.306 |
| *Beta macrocarpa* | 9 | 2.98 | -0.00474 | 0.296 | 0.000689 | -0.977 |
| *Centaurium tenuiflorum* | 1 | -1.01 | 0.00271 | 0.219 | 0.000576 | -0.422 |
| *Centaurium tenuiflorum* | 2 | -1.01 | 0.00271 | 0.219 | 0.000576 | -0.961 |
| *Centaurium tenuiflorum* | 3 | -1.01 | 0.00271 | 0.219 | 0.000576 | -0.441 |
| *Centaurium tenuiflorum* | 4 | -1.01 | 0.00271 | 0.219 | 0.000576 | 0.496 |
| *Centaurium tenuiflorum* | 5 | -1.01 | 0.00271 | 0.219 | 0.000576 | -0.121 |
| *Centaurium tenuiflorum* | 6 | -1.01 | 0.00271 | 0.219 | 0.000576 | 0.456 |
| *Centaurium tenuiflorum* | 7 | -1.01 | 0.00271 | 0.219 | 0.000576 | 0.342 |
| *Centaurium tenuiflorum* | 8 | -1.01 | 0.00271 | 0.219 | 0.000576 | 0.0909 |
| *Centaurium tenuiflorum* | 9 | -1.01 | 0.00271 | 0.219 | 0.000576 | 0.56 |
| *Hordeum marinum* | 1 | -0.714 | 0.00329 | 0.387 | 0.000693 | -0.999 |
| *Hordeum marinum* | 2 | -0.714 | 0.00329 | 0.387 | 0.000693 | -0.989 |
| *Hordeum marinum* | 3 | -0.714 | 0.00329 | 0.387 | 0.000693 | 1.51 |
| *Hordeum marinum* | 4 | -0.714 | 0.00329 | 0.387 | 0.000693 | 0.391 |
| *Hordeum marinum* | 5 | -0.714 | 0.00329 | 0.387 | 0.000693 | 0.724 |
| *Hordeum marinum* | 6 | -0.714 | 0.00329 | 0.387 | 0.000693 | 0.949 |
| *Hordeum marinum* | 7 | -0.714 | 0.00329 | 0.387 | 0.000693 | -0.965 |
| *Hordeum marinum* | 8 | -0.714 | 0.00329 | 0.387 | 0.000693 | -0.221 |
| *Hordeum marinum* | 9 | -0.714 | 0.00329 | 0.387 | 0.000693 | -0.398 |
| *Leontodon maroccanus* | 1 | -0.546 | 0.00338 | 0.536 | 0.000996 | 2.55 |
| *Leontodon maroccanus* | 2 | -0.546 | 0.00338 | 0.536 | 0.000996 | 2.05 |
| *Leontodon maroccanus* | 3 | -0.546 | 0.00338 | 0.536 | 0.000996 | -0.719 |
| *Leontodon maroccanus* | 4 | -0.546 | 0.00338 | 0.536 | 0.000996 | 0.213 |
| *Leontodon maroccanus* | 5 | -0.546 | 0.00338 | 0.536 | 0.000996 | -2.31 |
| *Leontodon maroccanus* | 6 | -0.546 | 0.00338 | 0.536 | 0.000996 | -0.397 |
| *Leontodon maroccanus* | 7 | -0.546 | 0.00338 | 0.536 | 0.000996 | -0.081 |
| *Leontodon maroccanus* | 8 | -0.546 | 0.00338 | 0.536 | 0.000996 | -0.404 |
| *Leontodon maroccanus* | 9 | -0.546 | 0.00338 | 0.536 | 0.000996 | -0.893 |
| *Parapholis incurva* | 1 | -0.522 | 0.00251 | 0.234 | 0.000701 | -1.05 |
| *Parapholis incurva* | 2 | -0.522 | 0.00251 | 0.234 | 0.000701 | 0.00531 |
| *Parapholis incurva* | 3 | -0.522 | 0.00251 | 0.234 | 0.000701 | -0.334 |
| *Parapholis incurva* | 4 | -0.522 | 0.00251 | 0.234 | 0.000701 | 0.433 |
| *Parapholis incurva* | 5 | -0.522 | 0.00251 | 0.234 | 0.000701 | 0.199 |
| *Parapholis incurva* | 6 | -0.522 | 0.00251 | 0.234 | 0.000701 | -0.189 |
| *Parapholis incurva* | 7 | -0.522 | 0.00251 | 0.234 | 0.000701 | 0.00271 |
| *Parapholis incurva* | 8 | -0.522 | 0.00251 | 0.234 | 0.000701 | 0.308 |
| *Parapholis incurva* | 9 | -0.522 | 0.00251 | 0.234 | 0.000701 | 0.628 |
| *Plantago coronopus* | 1 | 0.046 | 6.72e-05 | 0.16 | 0.000477 | 0.453 |
| *Plantago coronopus* | 2 | 0.046 | 6.72e-05 | 0.16 | 0.000477 | 0.485 |
| *Plantago coronopus* | 3 | 0.046 | 6.72e-05 | 0.16 | 0.000477 | -0.0366 |

| | | | | | | |
|---|---|---|---|---|---|---|
| *Plantago coronopus* | 4 | 0.046 | 6.72e-05 | 0.16 | 0.000477 | 0.00253 |
| *Plantago coronopus* | 5 | 0.046 | 6.72e-05 | 0.16 | 0.000477 | 0.0559 |
| *Plantago coronopus* | 6 | 0.046 | 6.72e-05 | 0.16 | 0.000477 | -0.615 |
| *Plantago coronopus* | 7 | 0.046 | 6.72e-05 | 0.16 | 0.000477 | -0.153 |
| *Plantago coronopus* | 8 | 0.046 | 6.72e-05 | 0.16 | 0.000477 | -0.275 |
| *Plantago coronopus* | 9 | 0.046 | 6.72e-05 | 0.16 | 0.000477 | 0.0824 |
| *Polypogon maritimus* | 1 | -1.63 | 0.00642 | 0.47 | 0.001 | -1.001 |
| *Polypogon maritimus* | 2 | -1.63 | 0.00642 | 0.47 | 0.001 | -0.792 |
| *Polypogon maritimus* | 3 | -1.63 | 0.00642 | 0.47 | 0.001 | -0.219 |
| *Polypogon maritimus* | 4 | -1.63 | 0.00642 | 0.47 | 0.001 | -1.098 |
| *Polypogon maritimus* | 5 | -1.63 | 0.00642 | 0.47 | 0.001 | 0.064 |
| *Polypogon maritimus* | 6 | -1.63 | 0.00642 | 0.47 | 0.001 | -0.582 |
| *Polypogon maritimus* | 7 | -1.63 | 0.00642 | 0.47 | 0.001 | 0.867 |
| *Polypogon maritimus* | 8 | -1.63 | 0.00642 | 0.47 | 0.001 | 2.999 |
| *Polypogon maritimus* | 9 | -1.63 | 0.00642 | 0.47 | 0.001 | -0.230 |

### 4 Posterior distributions and temporal evolutions

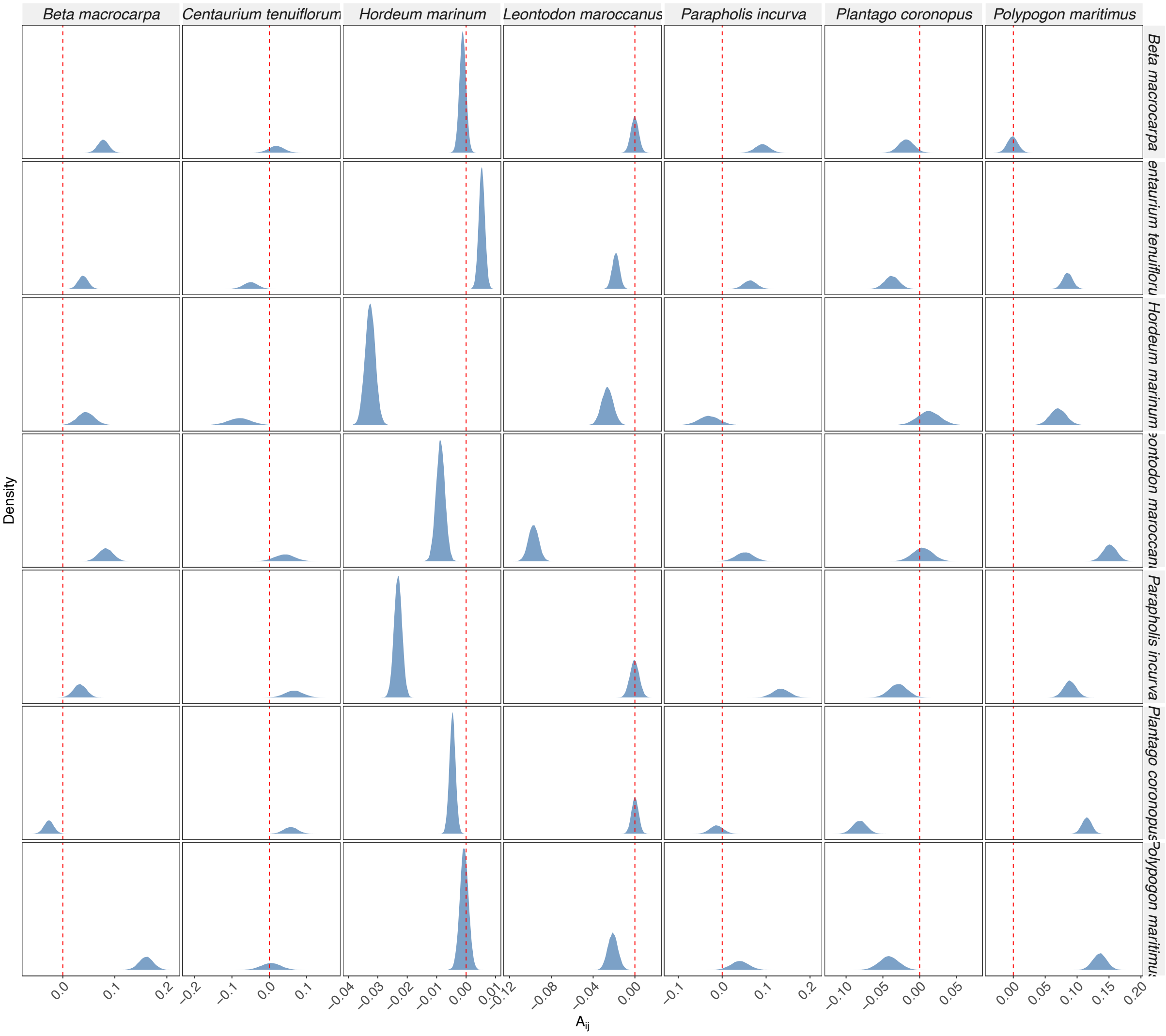


Figure 18: Posterior distribution of the values of the baseline interaction term $(A_{ij})$ for each species pair.

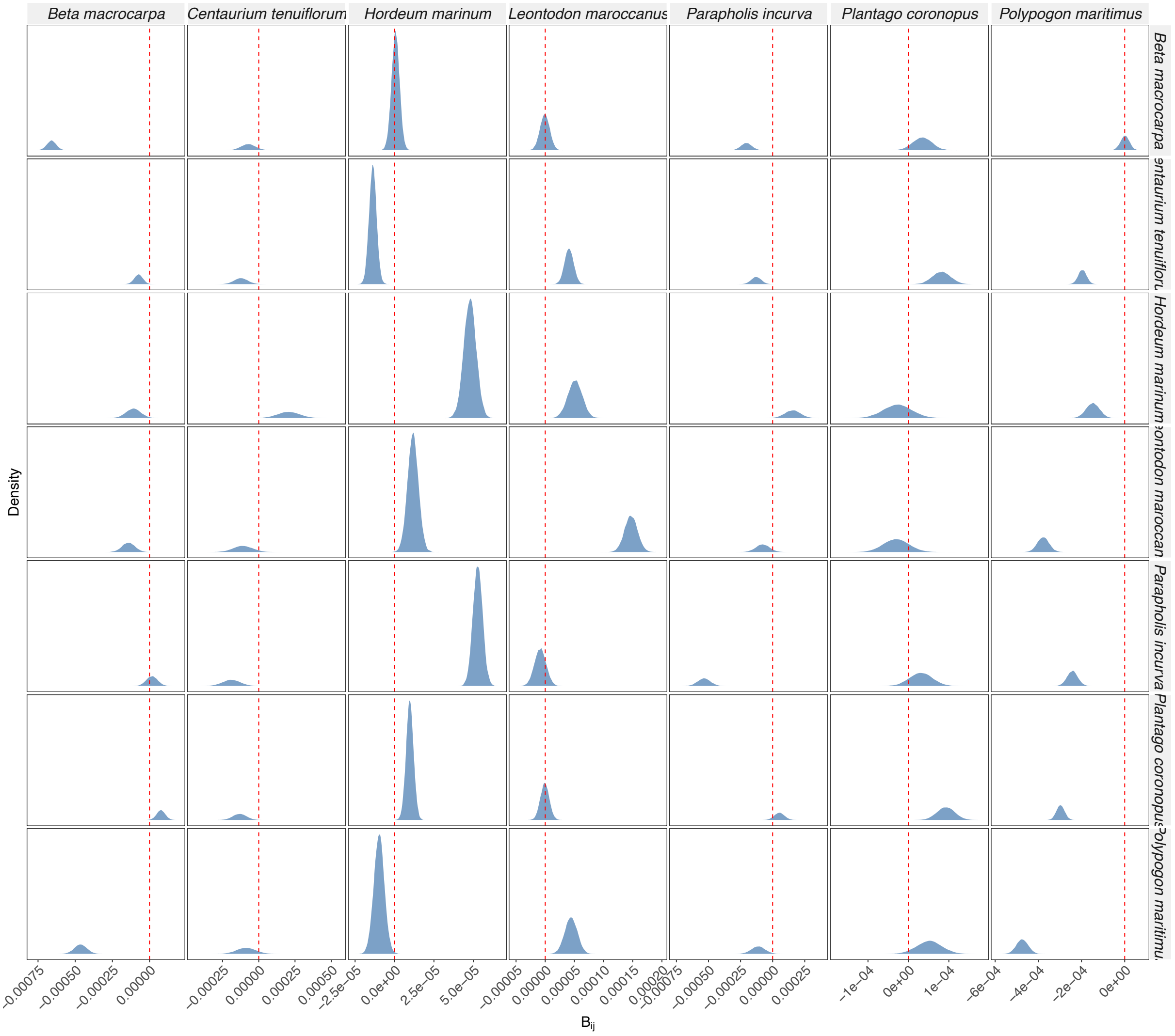


Figure 19: Posterior distribution of the effect of the time-varying environmental driver on interaction terms ($B_{ij}$) for each species pair.

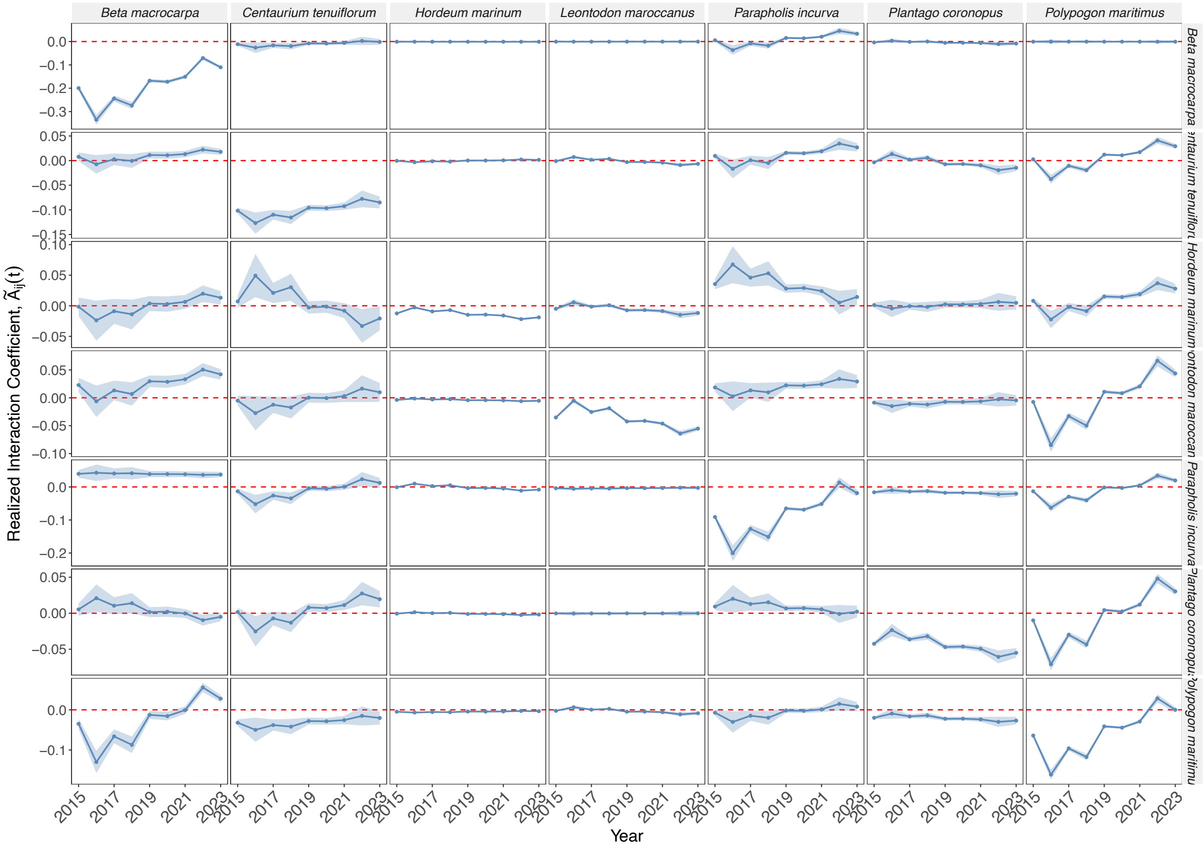


Figure 20: Temporal dynamics of realized interactions: $\tilde{A}_{ij}(t) = A_{ij} + B_{ij}\theta(t)$. Shaded regions indicate 95% Bayesian credible intervals.

### Time-varying interactions change coexistence mechanisms

Violeta Calleja-Solanas[1,2], Ignasi Bartomeus[2] and Oscar Godoy[2]
[1]Department of Biology, University of Oxford, Oxford, United Kingdom
[2]Estación Biológica de Doñana (EBD-CSIC), Americo Vespucio 26 41092, Seville, Spain

**Short running title:** Time-varying interactions change coexistence mechanisms

**Statement of authorship:** All authors conceptualized the ideas of the study. VC-S performed the data analysis. VC-S and OG drafted the manuscript with contributions from IB. OG provided the annual plant dataset.

**Data and code accessibility statement:** Datasets available at https://doi.org/10.5281/zenodo.7527011. Historical rainfall data in the area (Aznalcazar station) was shared by Meteoblue (for the 85-year average) and Junta de Andalucía. The code needed to reproduce this study can be downloaded from Zenodo https://zenodo.org/records/15625585.

**Keywords** : temporal networks | time-series data | coexistence metrics | environmental variability | ecological interactions | structural stability

- Type of article: Letter
- Word counts: Abstract, 139 words; main text, 4902 words
- This article includes 5 Figures, 20 Supplementary Figures, 4 Supplementary Tables, and 54 references.
- Mails of all authors:

Thanks to this unique dataset, we obtained empirical estimates of intrinsic growth rates $\tilde{r}_{t,b,i}$ that are variable in time $t$ (across 9 years) plus a stochastic spatial effect across the different sampling sites

$b$, and interactions $\tilde{A}_{t,ij}$ variable in time. The species belong to disparate taxonomic families and have different functional profiles (Table 1), but are primarily subjected to the same environmental driver, that is, annual rainfall. Moreover, they have recently experienced drought periods (Fig. S2), making them suitable for studying the consequences of droughts on community structure. The anomalies to study the effect of rainfall on stability and community composition are calculated as the difference from the long-term 85-year average (520 mm).

#### 2.2 Time-varying framework

Our theoretical approach is based on the Ricker model, a discrete version of the generalized Lotka-Volterra equations (Levine and HilleRisLambers, 2009). In its classic, deterministic form, changes in the abundance of species $i$ at time $t$ ($N_t^i$) are described by:

$$\log \frac{N_{t+1}^i}{N_t^i} = r_i + \sum_{j=1}^{n} A_{ij} N_t^j \tag{1}$$

where $r_i$ is the intrinsic growth rate of species $i$ and $A$ is the interaction matrix of the $n$ species. This baseline model does not depend on any temporal variable. The Ricker model is commonly applied to plant communities (Mayfield and Stouffer, 2017) because it can naturally accommodate both positive and negative interactions with varying strength, something that other standard population models such as the Beverton-Holt cannot do.

To explicitly link community dynamics to environmental fluctuations, we introduced time-varying species responses in intrinsic growth rates $r_i'$ and interactions $B_{ij}$ via effects of environmental conditions at each time $\theta_t$. In our system, the primary environmental driver is rainfall $\theta_t$. Furthermore, to account for local contingencies across our different sampling sites $b$, we incorporated hierarchical, site-level stochastic effects on species performances, $u_{i,b}$. Our proposed time-varying framework reads:

$$\log \frac{N_{t+1}^{i,b}}{N_t^{i,b}} = r_i + r_i'\theta_t + \sum_{j=1}^{n} (A_{ij} + B_{ij}\theta_t) N_t^{j,b} + u_{i,b}. \tag{2}$$

The time-varying terms of Eq. (2) involve a deterministic term on the growth rates $r_i'\theta_t$ and interactions $B_{ij}\theta_t$. By fitting this full model via Bayesian inference, we obtained joint posterior distributions for all baseline parameters and their environmental modifiers. From these posteriors, we calculated the distributions of the realized interaction matrices ($\tilde{A}_{t,ij}$) and realized intrinsic growth rates ($\tilde{r}_{t,b,i}$) for any given rainfall condition and site:

$$\tilde{A}_{t,ij} = A_{ij} + B_{ij}\theta_t, \tag{3}$$

$$\tilde{r}_{t,b,i} = r_i + u_{i,b} + r_i'\theta_t. \tag{4}$$

Hence, species interactions change annually in response to an environmental variable, and species exhibit different performance depending on that environmental driver and the site $b$ where they are sampled.

To study the long-term behavior of the system, in which the average per capita growth rates are all zero, we time-averaged the multispecies Ricker model. Assuming the populations persist over a sufficiently long time, the abundance change (left-hand side of Eq. (2)) tends to zero after some transient:

$$\frac{1}{T}\sum_{t}^{T} \log \frac{N_{t+1}^{i,b}}{N_t^{i,b}} = \frac{\log N_{T+1}^{i,b} - \log N_1^{i,b}}{T} \approx 0. \tag{5}$$

Applying the time-averaging operator $\frac{1}{T}\sum_t^T$, and denoting the time average of a variable $x_t$ as $\overline{x}$, we can map into the classic linear form $\vec{r} = -\mathbf{A}\vec{N}^*$, by defining the effective time-averaged interaction matrix elements $\overline{\tilde{A}_{ij}}$, Eq. (6), and the effective time-averaged intrinsic growth rates, Eq. (7):

$$\overline{\tilde{A}_{ij}} = A_{ij} + B_{ij}\overline{\theta}, \quad (6)$$

$$\overline{\tilde{r}_i} = r_i + r_i'\overline{\theta} + u_{i,b} + \sum_{j=1}^{n} B_{ij}\text{cov}(\theta, N^{j,b}). \quad (7)$$

The covariance term appears only in Eq. (7). It accounts for the effect of averaging a non-linear function over the temporal sequences of environmental conditions $\{\theta_1, \theta_2, \ldots, \theta_t, \ldots, \theta_T\}$ and population sizes $N$. As a consequence, two different sequences of environmental conditions with the same mean $\overline{\theta}$, such as a constant environment ($\theta_1 = \theta_2 = \cdots = \theta_T, \forall t$) and an oscillating one ($\{\theta_1, \theta_2, \theta_1, \theta_2, \ldots\}$), have the same $\overline{\tilde{A}}$, but differ in their covariance and then in their $\overline{\tilde{r}}$ (Supplementary Information, SI 1).

In a time-varying framework, both the shape and size of the feasibility domain may vary since they are dictated by $\tilde{A}$ (Allen-Perkins et al., 2023; Grilli et al., 2017), which may reorganize with rainfall. The position of $\tilde{r}$ may also change over time, as shown in Eq. (4). Because we estimated these parameters using Bayesian inference, we fully propagated this parameter uncertainty by calculating all structural stability metrics across the posterior draws. For the time-averaged approach, the effective time-averaged interaction matrix $\overline{\tilde{A}}$ only depends on the mean environmental conditions ($\overline{\theta}$), so two different environmental trajectories will have the same feasibility domain (size and shape) if they have a common mean environment ($\overline{\theta}$). On the contrary, time-averaged intrinsic growths $\overline{\tilde{r}_i}$ not only depend on $\overline{\theta}$ but also on a fluctuation-dependent term, the covariance between rainfall and the time series of species abundances. This covariance encodes the differences that arise from two different environmental trajectories. Then, time-varying environmental conditions do not change coexistence opportunities, but can alter the intrinsic growth rate vector across the feasibility domain via the covariance term.

#### 2.6 Simulations

We numerically explored how different environmental conditions affect the coexistence of the communities by accounting for different rainfall scenarios with the same mean $\overline{\theta} = 500$ mm: constant rainfall similar to the 85-year average, $\theta_{\text{cte}} = 500$ mm, decreasing rainfall $\theta_{\downarrow}$ from 600 to 400 mm, increasing rainfall $\theta_{\uparrow}$ from 400 to 600 mm, a triangular wave $\theta_{\wedge}$ alternating between 600 and 400 mm, and a sinusoidal sequence $\theta_{\sim}$ centered at 500 mm with 100 mm of amplitude and a 5-year period. To run these simulations, we drew 9000 parameter values from our posterior distributions and parameterized the multispecies Ricker equations for each draw until they reached (quasi)stationarity. We then recorded which parameter combinations led to all species persisting. We performed these simulations to obtain a computational approximation of the size of the feasibility domain. Then, we calculated the time-averaged $\overline{\tilde{A}}$ and $\overline{\tilde{r}}$ (neglecting the transient), Eqs. (6) and (7), to compare their feasibility predictions with the simulations.

Since we have inferred the parameters of the time-varying model, Eq. (2), we can also simulate how biodiversity unfolds in the short term, during our 9 year sampling period, under hypothetical rainfall regimes, by producing abundance time series for a given rainfall sequence $\theta = \{\theta_1, \ldots, \theta_T\}$, such as: constant rainfall equal to the 85-year average $\langle\theta\rangle = 520$ mm, oscillations between the minimum, $\min(\theta) = 225$ mm, and maximum, $\max(\theta) = 625.5$ mm, recorded rainfalls in our study period, increasing rainfall from $\langle\theta\rangle$ to $\max(\theta) + \sigma(\theta) \simeq 700$ mm, and decreasing rainfall and drought from $\langle\theta\rangle$ to $\min(\theta) - \sigma(\theta) \simeq 100$mm. To fully propagate the parameter uncertainty, we generated an ensemble of simulations for each scenario by sampling from the joint posterior distributions. The simulations were run for the same duration as our empirical dataset (9 timesteps). To track biodiversity changes, we calculated the Shannon index at each timestep $t$ and site $b$ across the simulation ensembles: $-\sum_i \left[\frac{N_t^{i,b}}{\sum_j N_t^{j,b}} \log\left(\frac{N_t^{i,b}}{\sum_j N_t^{j,b}}\right)\right]$. Ultimately, these simulations have three possible outcomes: (i) the presence of a persistent group of species (yielding a positive Shannon diversity), (ii) the complete species extinction of the community, or (iii) the dominance by a single species (in the last two cases, Shannon diversity is zero).

Simulations accounting for long-term dynamics with a $\theta_\wedge$ scenario (triangular fluctuations in rainfall) obtained the most feasible communities at the end of our simulations ($1762/9000 \approx 20\%$) than any other scenario, followed by constant rainfall $\theta_{\text{cte}}$ ($1534 \approx 17\%$) and sinusoid fluctuations in rainfall $\theta_\sim$ ($1471 \approx 16\%$). The decreasing and increasing rainfall scenarios had only 881 ($\sim 1\%$) and 234 ($\sim 0.3\%$) feasible communities, respectively. This means that the oscillating scenarios have among the largest numerically calculated feasibility domains. Interestingly, from these feasible communities, the time-averaged intrinsic growth rate overestimates and underestimates the feasibility of some scenarios depending on the sites (Fig. S8).

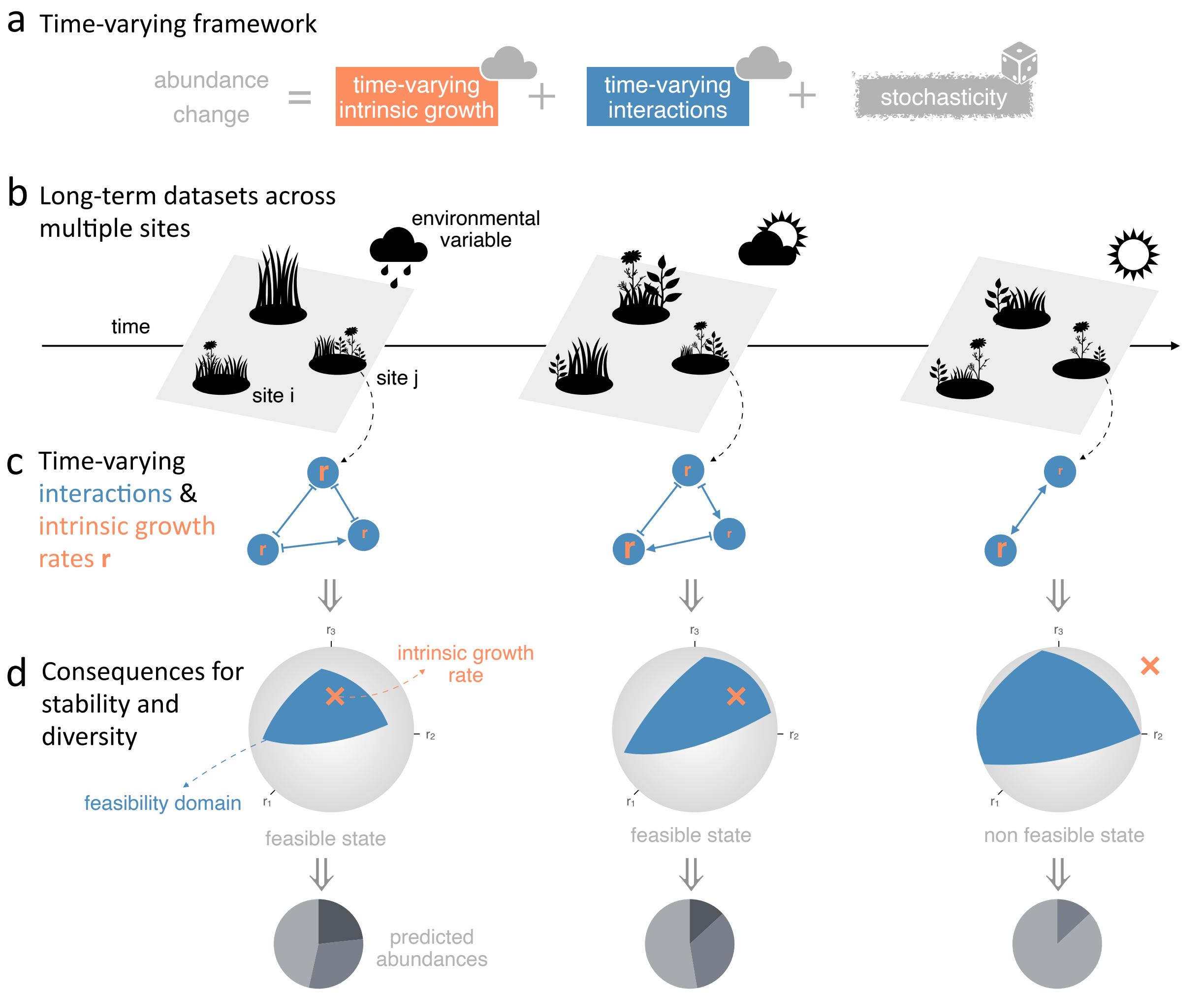


Figure 1: **Illustration of the theoretical framework in a toy community of 3 species.** (a) Estimation of interactions and intrinsic growth rates that vary through time according to changes in environmental factors. For each dataset, (b) thanks to well-resolved empirical observations from multiple sites constrained to rainfall, we can disentangle the time-varying and stochastic contributions of species' parameters to community dynamics, (c) obtaining time-varying species interactions $\tilde{A}$ (represented as blue networks) and intrinsic growth rates $\tilde{r}$ for each site. (d) In turn, these changes in species parameters modify the emerging properties of ecological communities, such as their stability (represented as the position of $\tilde{r}$, orange cross, within the feasibility domain, blue shape) and diversity.

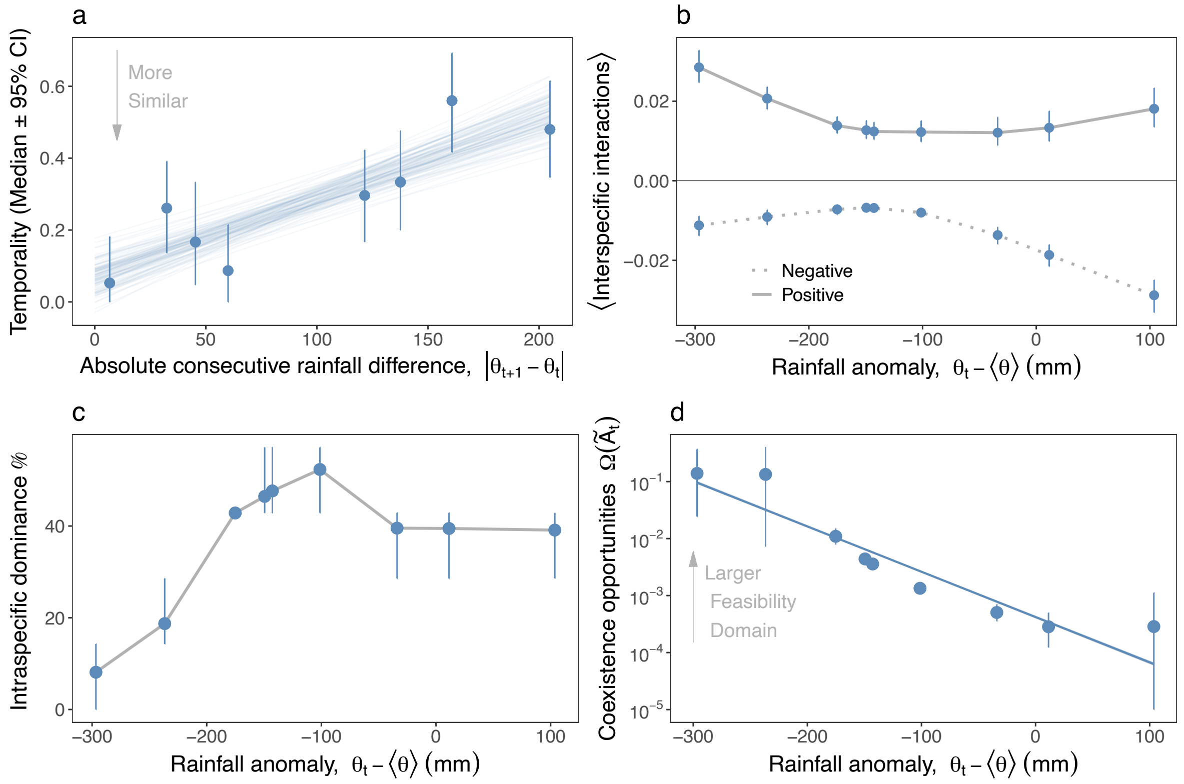


Figure 2: **Structural changes in interaction networks over time.** **a** Temporality, the degree of turnover of two consecutive interaction matrices. Blue solid lines are least-squares fits obtained separately within each of 100 randomly chosen posterior draws, so that their spread reflects posterior uncertainty in the interactions. **b** Mean interspecific positive (solid) and negative (dotted) interactions over species versus rainfall anomalies. **c** Changes in the percentage of species whose self-regulation dominates their interspecific interactions with rainfall anomalies. Smaller values indicate less self-regulation. Gray lines for visualization guidance. **d** Coexistence opportunities (size of the feasibility domain for each realized interaction matrix $\Omega(\tilde{A}_t)$) with respect to rainfall anomalies. For each rainfall anomaly, properties of the interaction network are calculated with 1000 posterior draws. Solid blue lines are Bayesian generalized linear model fits for 100 draws. Note that gray lines in **b** and **c** are for visualization purposes. SI 4 contains the posterior distributions and temporal evolution of the values of the time-varying networks.

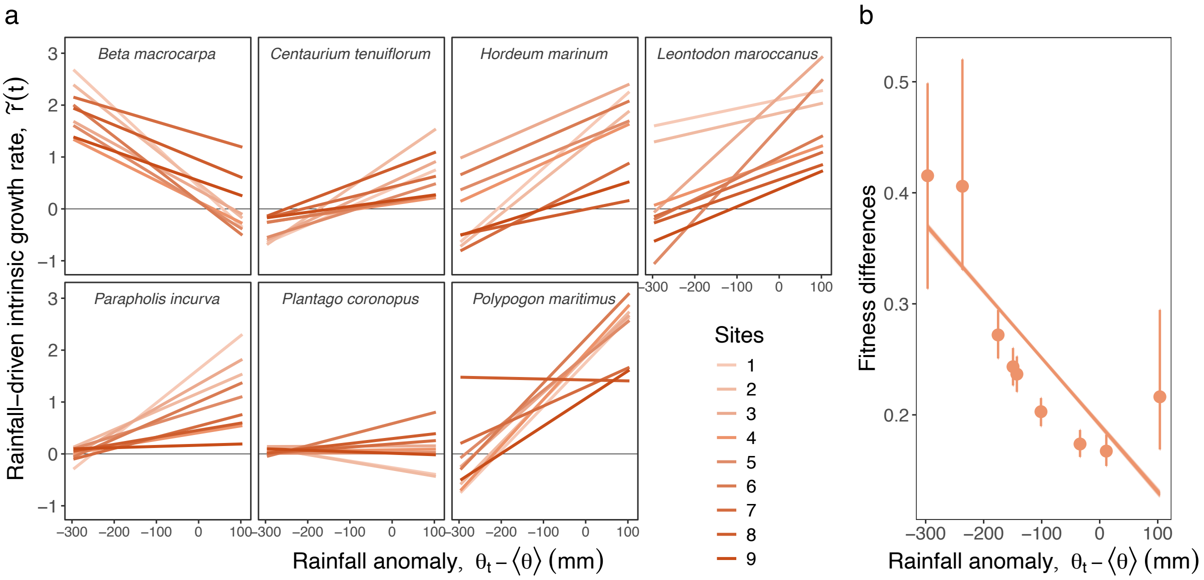


Figure 3: **Species' performance explained by deterministic effects of rainfall and spatial context dependency**. **a** Realized intrinsic growth rates for all sampled sites. A relationship parallel to the x-axis indicates no sensitivity to rainfall anomalies, and a greater slope variation for a species means more sensitivity to the site's conditions. Only the posterior distribution mean is plotted here. **b** Fitness differences associated with rainfall anomalies, measured as the distance between $\tilde{r}$ and the incenter. Each point is the average across sites of 1000 posteriors draws, and solid orange lines are Bayesian generalized linear model fits for 100 draws. SI 4 contains the temporal evolution of the values of the time-varying $\tilde{r}(t)$.

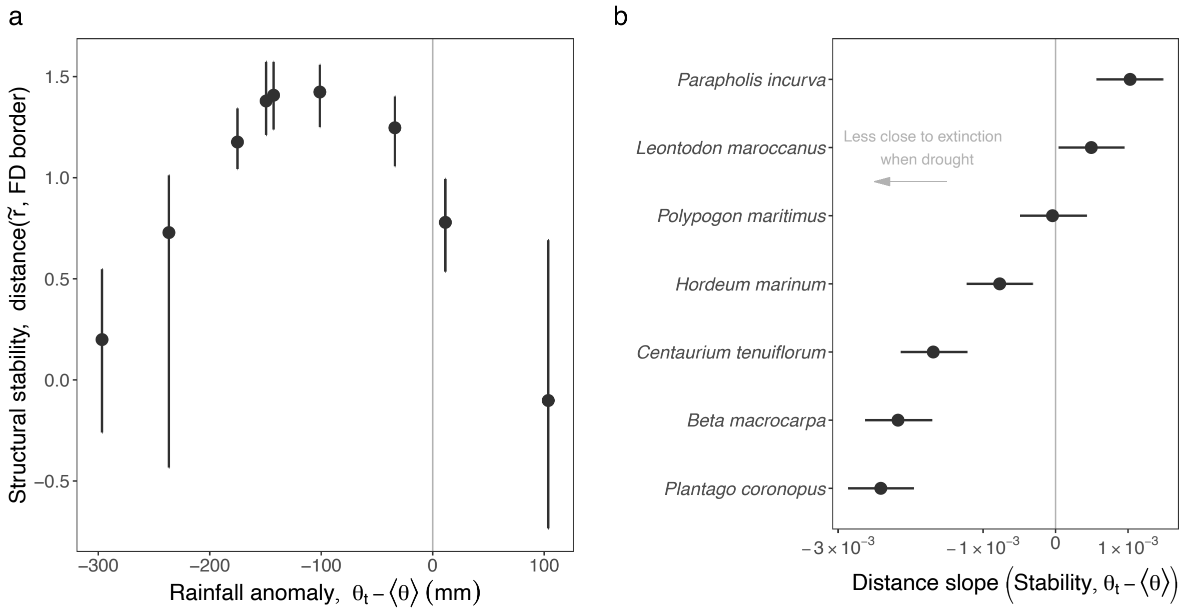


Figure 4: **Consequences on stability of temporal changes in coexistence mechanisms. a** Structural stability as the average over the whole community and sites of the species distance between intrinsic growth rates and the nearest border of the feasibility domain (FD) where a species goes extinct. **b** Slopes of the distance between intrinsic growth rates and the nearest border of the feasibility domain (FD) for each species as a function of rainfall anomalies. The trends for each species are in Fig. S7. In our sign convention, positive distances denote that a species is more prone to persist in wetter years, and negative slopes indicate greater species persistence in drier years. Error bars represent the standard deviations across all sites for 1000 draws each.

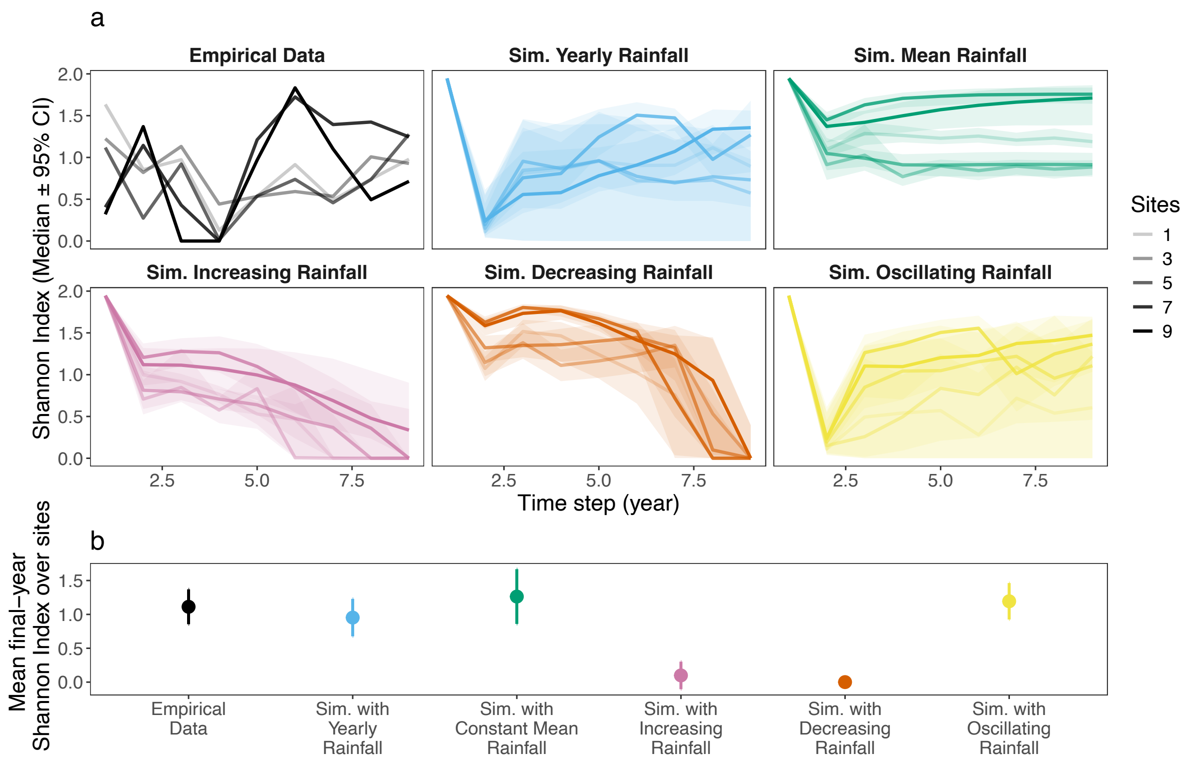


Figure 5: **Consequences on diversity of rainfall patterns**. **a** Shannon diversity index for different simulated rainfall regimes. Besides the index for the empirical abundances (black), the lines show 1000 simulations from our estimated model for different scenarios: rainfall varies according to the recorded values, rainfall does not vary and is equal to the average, increases over time, decreases, and oscillates between the maximum and minimum values over 9 years. **b** Shannon indices for the final state of the simulations. Since our community size is $n = 7$, the maximum value of the index is 2.

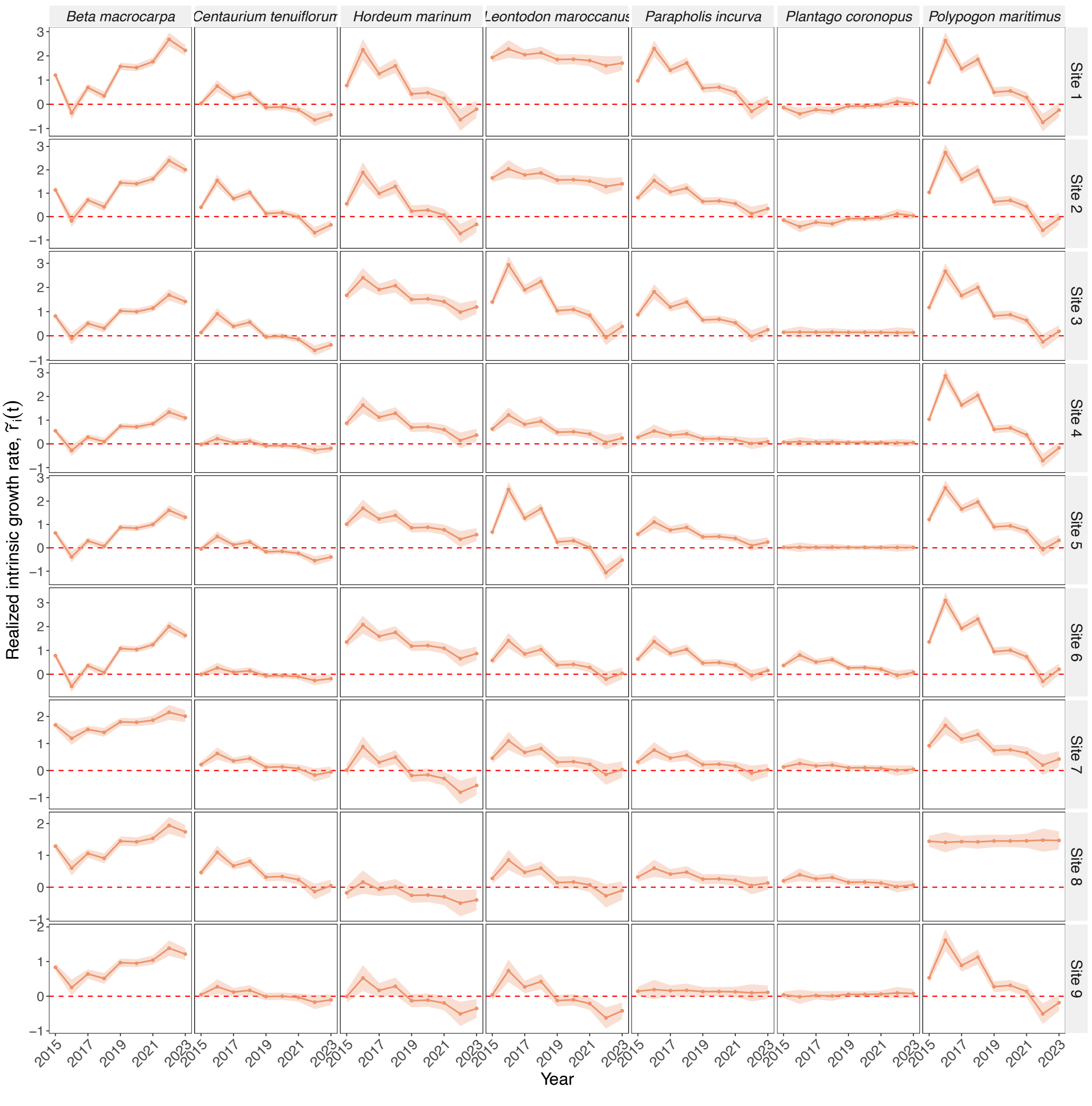


Figure 21: Temporal dynamics of realized intrinsic growth rates: $\tilde{r}(t)$. Rows: Sites. Columns: Species. Shaded regions indicate 95% Bayesian credible intervals.
